# Global pleiotropic effects in adaptively evolved *Escherichia coli* lacking CRP reveal molecular mechanisms that define growth physiology

**DOI:** 10.1101/2020.06.18.159491

**Authors:** Ankita Pal, Mahesh S. Iyer, Sumana Srinivasan, Aswin S Seshasayee, K.V. Venkatesh

## Abstract

Evolution entails the orchestration of cellular resources together with mutations to achieve fitter phenotypes. Here, we determined the system-wide pleiotropic effects that redress the significant perturbations caused by the deletion of global transcriptional regulator CRP in *Escherichia coli* when evolved in the presence of glucose. We elucidated that absence of CRP results in alterations in key metabolic pathways instrumental for the precise functioning of protein biosynthesis machinery that subsequently corroborated with intracellular metabolite profiles. Apart from acquiring mutations in the promoter of glucose transporter *ptsG*, the evolved populations recovered the metabolic pathways to their pre-perturbed state with amelioration of protein biosynthesis machinery coupled with fine-tuned proteome re-allocation that enabled growth recovery. However, ineffective utilization of carbon towards biomass as perceived from ATP maintenance flux and costly amino acid accumulations poses a limitation. Overall, we comprehensively illustrate the genetic and metabolic adjustments underlying adaptive evolvability, fundamental for understanding the growth physiology.

## Introduction

Global transcriptional factors (TFs), represent a cornerstone in the transcriptional regulatory network (TRN), that facilitates system-wide changes in gene expression levels in response to alterations in its external or internal environment (Babu et al., 2004; Martínez-Antonio and Collado-Vides, 2003; Seshasayee et al., 2011). Considering the complex interactions existing within the TRN of an organism, the absence of global TFs results in direct or indirect cellular responses that incapacitate the ability to attain favourable phenotypic outcomes, even for a simple prokaryote like *E. coli*. Regulatory mechanisms of CRP (cAMP receptor protein), a global transcriptional regulator in *E. coli*, under diverse conditions has been an area of research over many decades. CRP, along with its cognate signalling molecule cAMP, activates transcription at more than 200 promoters as evidenced from the genome-wide binding studies (Hollands et al., 2007; Kolb et al., 1993). *In vitro* and *in vivo* binding assays have predicted and validated such genome-wide binding sites of CRP, wherein, its interactions with RNA polymerase at such promoters have been addressed (Grainger et al., 2005; Latif et al., 2018). Specifically, CRP has been shown to regulate, i) genes involved in the transport and metabolism of glucose and other carbon sources like mannose and galactose (Franchini et al., 2015; Gosset et al., 2004; Kim et al., 2018; Shimada et al., 2011), ii) genes encoding enzymes of the TCA cycle and oxidative phosphorylation (Kim et al., 2018; Nam et al., 2005; Zheng et al., 2004), and iii) genes of the stress response (Castanie-Cornet and Foster, 2001) under diverse nutritional conditions by classical approaches and microarray-based studies. Moreover, the physiological significance of its coactivator cAMP, in coordinating the carbon and nitrogen demands via carbon catabolites addressed the long-standing debate on carbon catabolite repression (You et al., 2013).

Despite the huge repository of data available for CRP, several questions are still unanswered. Since, changes in glucose uptake rate can only in part explain the changes in growth rate, the underlying molecular mechanisms driven by CRP, which determine the detrimental shift in exponential growth profiles, remain obscure. Scarce knowledge of the fate of cAMP and downstream metabolite signatures propagated by gene expression changes in the absence of CRP limits our ability to link molecular consequences to growth. Further, in the absence of CRP, proteome allocation principles (O’Brien et al., 2016; Scott et al., 2010; You et al., 2013) that facilitate system-wide fine-tuning of the necessary and unnecessary proteome towards biomass synthesis, remains unexplored. Thus, a fundamental question that arises now is, whether these represent the major drivers that facilitate a system-level response to ensure growth fitness in the event of the evolution of such a dysregulated mutant.

By exploiting such characteristics, we sought to investigate how an *E. coli* K-12 MG1655 strain lacking CRP, can cope up with this global disruption using adaptive laboratory evolution (ALE) under glucose minimal media conditions. ALE entails orchestration of genetic as well as phenotypic behaviour in response to mutations that provide growth fitness benefits to organisms under strict selection pressures (LaCroix et al., 2015; Mozhayskiy and Tagkopoulos, 2013). While a majority of the ALEs have focussed on understanding the adaptive rewiring in response to the loss of metabolic genes (Charusanti et al., 2010; McCloskey et al., 2018a; Notley-McRobb and Ferenci, 2000), studies that focus on ALEs on the loss of global transcriptional regulators, are now emerging (Lamrabet et al., 2019; Srinivasan et al., 2015). A previous study on *E. coli* REL606 and its evolved clones lacking CRP had elucidated how the organism copes with the growth defect by accumulating mutations in the promoter of the glucose transporter gene, *ptsG* (Lamrabet et al., 2019). Their work highlighted that generation of new transcription start sites in the promoter of *ptsG* and *manX* resulted in growth recovery. However, the global pleiotropic effects of such mutations on cellular processes that enabled growth recovery in the evolved populations has not been examined earlier. Therefore, to decipher the underlying molecular bases of divergence of evolved strains away from their ancestor (Dragosits et al., 2013; Ostrowski et al., 2005; Utrilla et al., 2016; Zorraquino et al., 2017), examining the evolution of a *crp* mutant with integration of transcriptomics, metabolomics and proteome allocation aspects can be a study of great value. Moreover, the nature of mutations and its molecular consequences are largely influenced by the genotype of the ancestral organism and growth phase wherein the serial passages are carried out (LaCroix et al., 2017; Rodŕiguez-Verdugo et al., 2016; Sandberg et al., 2014).

In this present study, using a multi-omics approach, we examine the global physiological significance of CRP for exponential growth in glucose minimal media conditions. We elucidate its underlying direct or indirect regulatory mechanisms that coordinate metabolite profiles and the allocation of proteomic resources along with its congruent gene expression change towards its cellular objectives. Further, we examined the pleiotropic effect of beneficial mutations on cellular processes underlying growth fitness in an *E. coli* K-12 MG1655 strain lacking CRP, under glucose minimal media conditions. Overall, by evaluating such genotype-phenotype relationships in the parent and evolved strains, we elucidate the inherent constraints of genetic and metabolic networks underlying evolvability in *E. coli*.

## Results

### Loss of CRP caused large shifts in the transcriptome of key metabolic pathways

CRP occupies an apex node in the hierarchical transcriptional regulatory network (TRN) of *E. coli* regulating a myriad of genes under diverse nutritional conditions (Martínez-Antonio and Collado-Vides, 2003). Here, we determined the role of CRP in regulating key metabolic pathways with glucose as a carbon source. Towards this, we first addressed the systemic effect caused by the loss of CRP by performing high-coverage RNA sequencing of Δ*crp* mutant in glucose minimal media condition during the mid-exponential phase. The transcriptome of this strain when compared to its parent wild-type (WT) strain, showed ~722 differentially expressed (DE) genes (Absolute Fold change (aFC) ≥ 2, Adjusted P-value (adj-P) < 0.05) of which ~544 genes (75%) were downregulated and ~ 178 genes (25%) were upregulated in the mutant (fig. 1A), indicating a large upset of the global transcriptome (supplementary file 1). This reiterated the role of CRP as a transcriptional activator, that was in good agreement with previous gene expression studies (Gosset et al., 2004; Latif et al., 2018; Zheng et al., 2004) (supplementary file 1).

**Figure 1.**
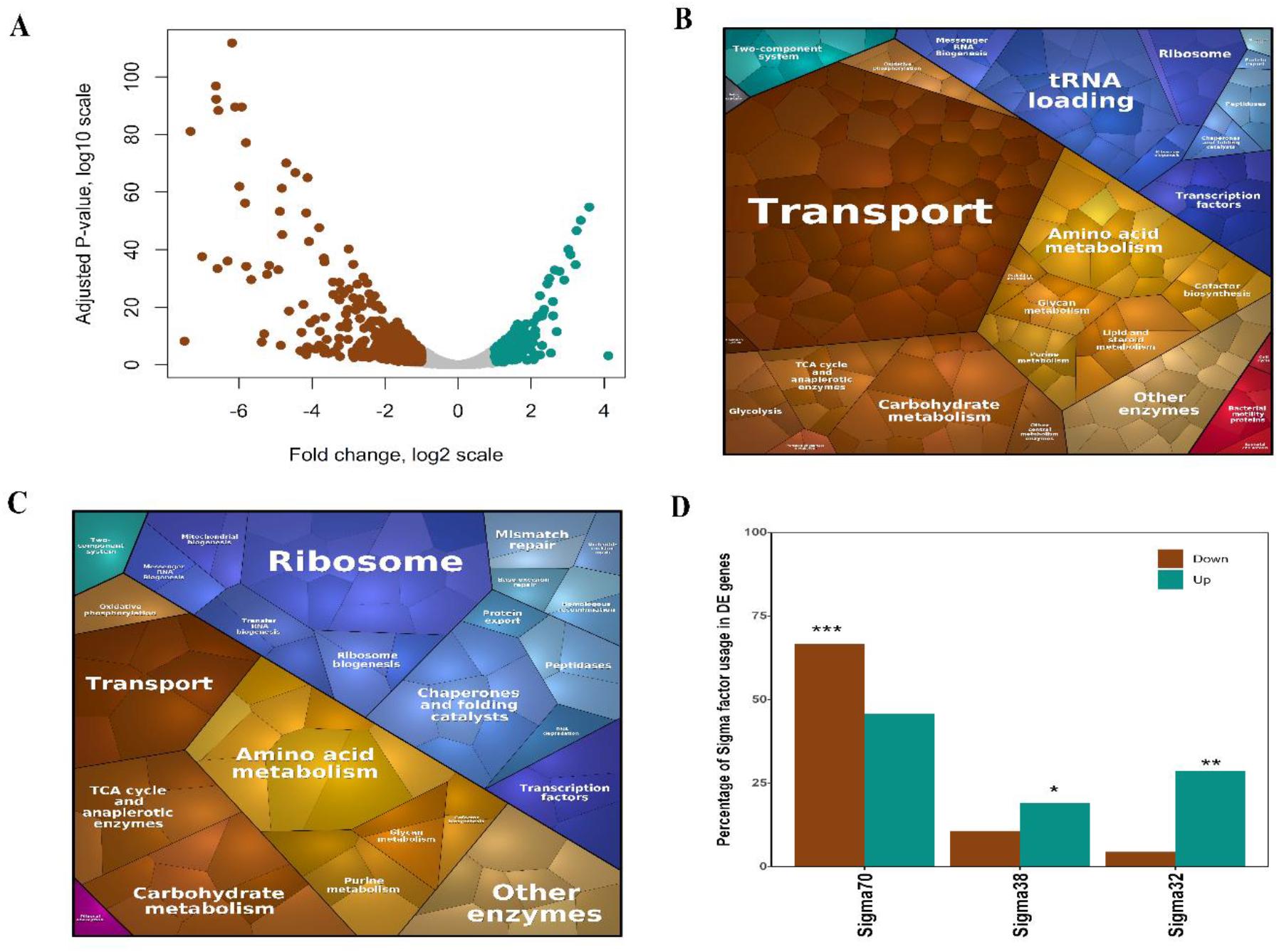
Severe perturbation of gene expression in Δ*crp* compared to WT. (A) Volcano plot of the DE genes Δ*crp* compared to WT depicted as Adj-P value (log10 scale) vs Fold change (log2 scale). The brown dots indicate downregulated genes and the cyan dots indicate upregulated genes. (B) The Voronoi treemaps showing the downregulated metabolic pathways enriched by KEGG classification. Transport (adj-P < 0.05), tRNA loading (adj-P < 10^−4^) and TCA cycle (adj-P < 0.05) were found to be significantly downregulated. (C) The Voronoi treemaps showing the upregulated metabolic pathways enriched by KEGG classification. Ribosome (adj-P < 10^−5^), Chaperone and folding catalysts (adj-P < 10^−2^) and TCA cycle (adj-P < 0.05) were found to be significantly upregulated. The genes enriched within each pathway are shown in supplementary fig. S1. Note: The size of the hexagon within each pathway is directly proportional to the absolute fold change observed for the genes. The colour of the hexagon denotes the specific pathways classified by KEGG. (D) Enrichment of genes under regulation of sigma factors; the brown bars and the cyan bars indicate the fraction of downregulated and upregulated genes in Δ*crp* vs to WT respectively. Significant increase or decrease is denoted by asterisks: one asterisk indicates P < 0.05, two asterisks indicate P < 10^−4^ and three asterisks indicate P < 10^−10^.

We examined the KEGG pathways that were significantly enriched among these DE genes and represented these pathways as Voronoi treemaps (fig. 1B-C and supplementary fig. S1). Our data indicated downregulated genes significantly associated with pathways such as transporters, tricarboxylic acid (TCA) cycle and anaplerotic enzymes needed for aerobic respiration, and tRNA loading pathways involved in for proper channelling of amino acids towards protein translation. Downregulation of transporters pathway was associated with the major glucose transporter *(ptsG)*, secondary glucose transporters *(manXYZ, malEFGKX*, and *lamB* that function under glucose limitation), transport of amino acids *(tdcC, proVXW, hisJ, livJKH*, *lysP*, *leuE*), and nucleotides (*tsx*, *uraA*, *nupCGX*), as well as alternate carbon transporters (*glpF*, *fruB*). Moreover, we observed the upregulated genes to be significantly associated with pathways such as ribosomes, and chaperone and folding catalysts. These are responsible for the general stress response of the cell, required to maintain proper protein turnover and integrity (Jozefczuk et al., 2010). Owing to the huge investment of proteomic resources for the synthesis of ribosomes (Scott et al., 2010; Zhu and Dai, 2019), such an increase in ribosome transcript levels might indicate an increase in unnecessary ribosomal proteins, thereby reducing the proteome share for metabolic proteins. Further, we found upregulated genes enriched in the TCA cycle and anaplerotic enzymes pathway to be primarily involved in glycolate and glyoxylate degradation (*glcDEF, aceA*), which might indicate an increase in hedging mechanism related to alternate carbon metabolism (Solopova et al., 2014; Utrilla et al., 2016). Overall, the considerable shifts in the transcriptome of these pathways, emphasize the metabolic dysregulation coupled to impaired functioning of protein biosynthesis machinery caused by the loss of a global regulator.

We identified a significant fraction of enriched downregulated genes (~35%, P < 10^−14^) that were found to be regulated by CRP as opposed to the upregulated genes (~9%, P > 0.1). Such gene expression patterns could be attributed to the direct effect of loss of CRP regulation as well as to indirect effects, mediated by global physiological factors (RNA polymerase, ribosome, and metabolites) (Berthoumieux et al., 2013; Klumpp et al., 2009) or ppGpp, the stringent response regulator (Traxler et al., 2008). However, studies have shown that upregulation in ribosomes could be independent of the effects of ppGpp due to a simple activation program driven by cellular demands (Scott et al., 2014). Additionally, ~66% (P < 10^−6^) of enriched downregulated genes, were found to be regulated by sigma 70, the major growth-related sigma factor associated with RNA Polymerase (fig. 1D), corroborating the association of CRP with sigma 70 reported previously (Grainger et al., 2005; Latif et al., 2018). A significant fraction of enriched upregulated genes was regulated by stress-related sigma factor, sigma 38 (~28%, P < 10^−2^) and sigma 32 (~22%, P < 10^−13^). Presumably, this suggested the reallocation of RNA Polymerase away from growth and towards stress-related genes as an indirect consequence of the loss of a global regulator.

### Adaptive evolution recapitulates mutations in the *ptsG* intergenic region but with altered binding profiles

To decipher how Δ*crp* copes with the perturbations in its global gene expression, we adaptively evolved five independent replicates of the mutant with multiple passages in batch culture with non-limiting glucose, strictly during the mid-exponential phase (supplementary fig. S2). We observed that the growth defect in Δ*crp* was rapidly recovered within ~100 generations of adaptive evolution (supplementary fig. S3) and the endpoint populations (EvoCrp) (Supplementary table S1) were further characterized in this study.

We performed whole-genome resequencing (WGS) to identify the causal mutations in all the EvoCrp endpoint populations (supplementary table S2). Out of eleven unique mutations detected across all the EvoCrp strains (described in greater detail in supplementary text), six were in the upstream promoter sequence of the *ptsG* gene, a component of the Phosphotransferase System (PTS) responsible for ATP-independent glucose uptake in *E. coli*. The intergenic mutations were mostly SNPs specifically in the operator sites of the repressors namely Fis (Cho et al., 2008; Shin et al., 2003), Mlc (Plumbridge, 2002; Plumbridge and Upr, 1998), and ArcA (Jeong et al., 2004), reported to repress*ptsG* gene expression (supplementary fig. S4). We, therefore, hypothesized that these mutations in the *ptsG* promoter region are responsible for altered binding affinity, resulting in the de-repression of the *ptsG* gene.

First, to examine the adaptive role of the mutations, we introduced the IG116 promoter mutation (as annotated in supplementary table S2) in the Δ*crp* background. The introduction of the mutation resulted in 85% recovery to the WT growth rate thereby confirming the adaptive nature of the promoter mutations in the EvoCrp strains (supplementary fig. S5A and S5B). Next, to test our altered binding hypothesis, we characterized the *in-vivo* binding affinity of Fis in IG116-Δ*crp* mutant strain using CHIP-qPCR. We did not observe any significant enrichment for Fis binding to the promoter neither in IG116 Δ*crp* strain nor in WT and Δ*crp* strain (supplementary fig. S6A-C), that was in agreement to a previous study in *E. coli* K-12 MG1655 (Cho et al., 2008). Also, as operator sequence encompassing IG116 mutation is also known to possess a binding affinity for Mlc (Plumbridge, 2002; Plumbridge and Upr, 1998), we characterized the binding affinity of Mlc in IG116 Δ*crp* strain, which also revealed similar results as in the case of Fis (supplementary fig. S6D). This emphasized the fact that the regulation of *ptsG* is not dependent on the interplay of the regulators Fis and Mlc. Besides, a detailed analysis of the WGS data indicated that most of the mutations in the *ptsG* promoter resulted in new transcriptional start sites (TSS) with “Pribnow” box-like consensus sequence, thereby reiterating that these *ptsG* promoter mutations augment the affinity of RNA Polymerase sigma 70 as previously observed in a recent study albeit in a different genetic background (Lamrabet et al., 2019). However, the mutation profile and the binding profile of negative regulators on the *ptsG* promoter in our study were markedly different from the previous study that can be attributed to the underlying differences in the genetic background of the parent strains used for evolution in both the studies.

### Adaptive rewiring of gene expression of metabolic pathways in EvoCrp strains

To understand the gene expression changes that enabled higher growth of the evolved strains, we first characterized the transcriptome of the EvoCrp strains by comparing it to Δ*crp* parent strain and identified the statistically significant DE genes (aFC ≥ 2, adj-P < 0.05) (supplementary file 1). Broadly, all EvoCrp strains compared to Δ*crp* showed fewer DE genes (ranging from ~75 to ~300 DE genes) as opposed to ~700 DE genes observed in Δ*crp* compared to WT (fig. 2A). A higher proportion of DE genes in EvoCrp showed an upregulation and a higher median expression level (P < 10^−6^, Mann Whitney test, fig. 2B and supplementary fig. S7 and S8) relative to Δ*crp*, implying a shift in transcriptome towards that align with increased growth. To understand the adaptive response of EvoCrp, we computed the correlation between the fold-change of gene expression in Δ*crp* vs WT and EvoCrp vs WT (fig. 2C and supplementary fig. S9) and observed a strong positive correlation (Pearson correlation coefficient, r = 0.74, P < 10^−15^). Also, the magnitude of the difference of fold change between EvoCrp and WT was lesser compared to Δ*crp* vs WT (P < 10^−15^, Mann Whitney test). Both of these observations indicated a partial restoration in EvoCrp gene expression states towards WT levels.

**Figure 2.**
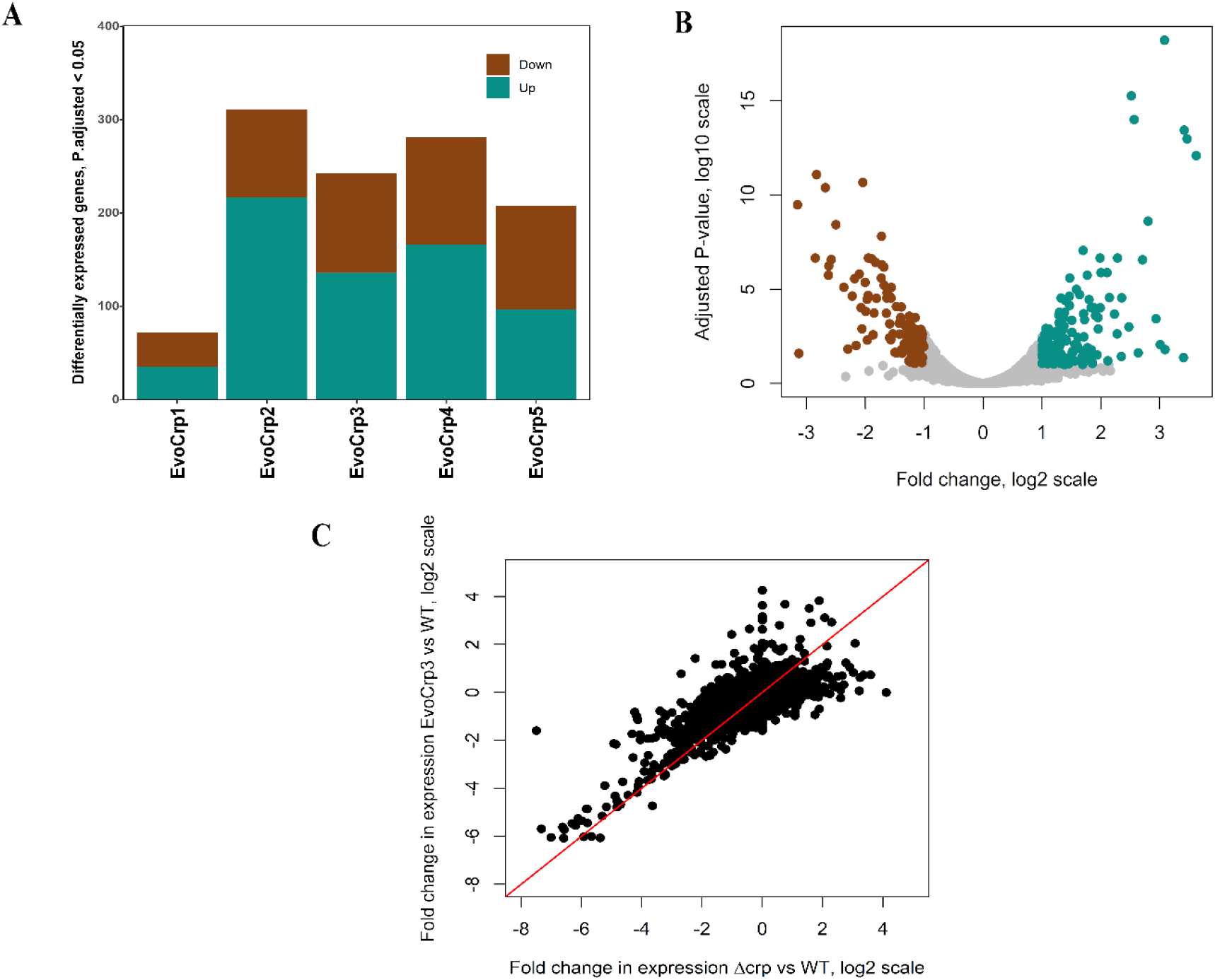
Transcriptome comparison of the EvoCrp compared to Δ*crp*. (A) Stacked plots showing the number of DE genes across all the EvoCrp strains. The downregulated genes are shown in brown and the upregulated genes are shown in cyan. (B) Volcano plot of the DE genes for one of the representative evolved strain, EvoCrp3 vs Δ*crp* depicted as Adj-P value (log10 scale) vs Fold change (log2 scale). See supplementary fig. S7 for volcano plots of DE genes for other EvoCrp strains. (C) A correlation plot between EvoCrp3 vs WT and Δ*crp* vs WT (Pearson correlation coefficient, r = 0.74, P < 10^−15^). See supplementary fig. S9 for correlation plots of DE genes for other EvoCrp strains.

Next, we investigated the metabolic pathways enrichment in EvoCrp strains. We observed that majority of the pathways that were affected in Δ*crp* were reverted in the EvoCrp strains (supplementary fig. S10) to the pre-perturbed state. For instance, across a majority of EvoCrp strains (fig. 3A-3B and supplementary fig. S11-S15), we observed upregulated genes significantly associated with transporters and tRNA loading pathways, and downregulated genes significantly associated with ribosome, and chaperone and folding catalyst related pathways. This indicated the restoration of protein turnover and folding machinery in EvoCrp strains. However, the number of genes belonging to the restored pathways agrees well with the partial restoration seen from the correlation plot (fig. 2C and supplementary fig. S10). The restoration in the gene expression in EvoCrp also suggested a reallocation of RNA polymerase, as indicated by the large fraction of the upregulated genes regulated by RNA polymerase sigma 70 (~76%, P < 10^−5^) and the downregulated genes regulated by stress-related sigma factor, sigma 38 (~37%, P < 10^−3^) and sigma 32 (~14%, P <10^−13^) (fig. 3C & supplementary fig. S16). Overall, in EvoCrp strains, we observed rewiring of metabolic pathways and amelioration of protein biosynthesis machinery, which corroborated with the reallocation of RNA polymerase favouring growth over stress-related functions.

**Figure 3.**
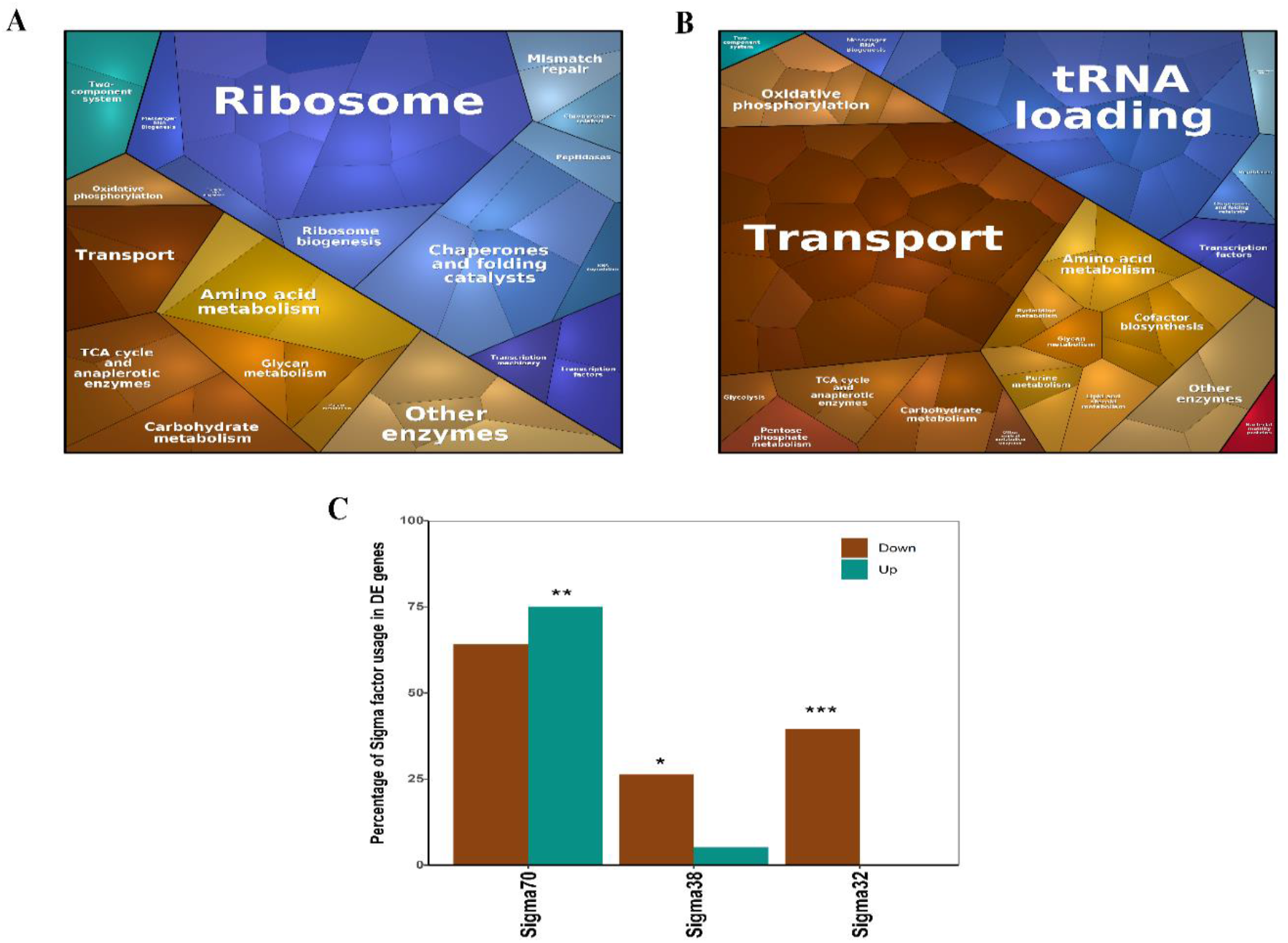
Enrichment analysis of the DE genes in EvoCrp3 compared to Δ*crp*. (A) The Voronoi maps showing the downregulated metabolic pathways enriched by KEGG classification. Ribosome (adj-P < 10^−8^), and Chaperone and folding catalysts (adj-P < 10^−2^) were found to be significantly downregulated. (B) The Voronoi treemaps showing the upregulated metabolic pathways enriched by KEGG classification. Transport (adj-P < 10^−3^), tRNA loading (adj-P < 10^−13^) and Oxidative phosphorylation (adj-P < 10^−5^) were found to be significantly upregulated. See supplementary fig. S11-S15 for the Voronoi maps of the pathways and the genes within each pathway for all EvoCrp strains. The genes enriched within each pathway are shown in supplementary fig. S13. Note: The size of the hexagon within each pathway is directly proportional to the absolute fold change observed for each gene. The colour of the hexagon denotes the specific pathways classified by KEGG. (C) Enrichment of genes under regulation of sigma factors; the brown bars and the cyan bars indicate the percentage of downregulated and upregulated genes in EvoCrp3 vs to Δ*crp* respectively. Significant increase or decrease is denoted by asterisks: one asterisk indicates P < 0.05, two asterisks indicate P < 10^−4^ and three asterisks indicate P < 10^−10^. See supplementary fig. S16 for sigma factor usage of DE genes for other EvoCrp strains.

### Accumulation of metabolites illustrate strain-specific growth effects

Next, to evaluate the metabolite levels that mirror the shifts in growth profiles (Barupal et al., 2013; Erickson et al., 2017), we characterized several metabolites of central carbon metabolism in the exponential phase of growth in Δ*crp* and the evolved strains using ^13^C-labelled metabolomics (supplementary file 2). An initial exploratory assessment was performed using partial least squares discriminatory analysis (PLS-DA) on 24 metabolites that were found to be statistically significant (adj-P < 0.05, Student’s t-test) in either EvoCrp vs Δ*crp* or EvoCrp vs WT or Δ*crp* vs WT strains. The two components of the PLS-DA scores plot (fig. 4A) which accounted for ~68% of the variance, revealed partial restoration of the metabolite profiles in EvoCrp towards WT levels.

**Figure 4.**
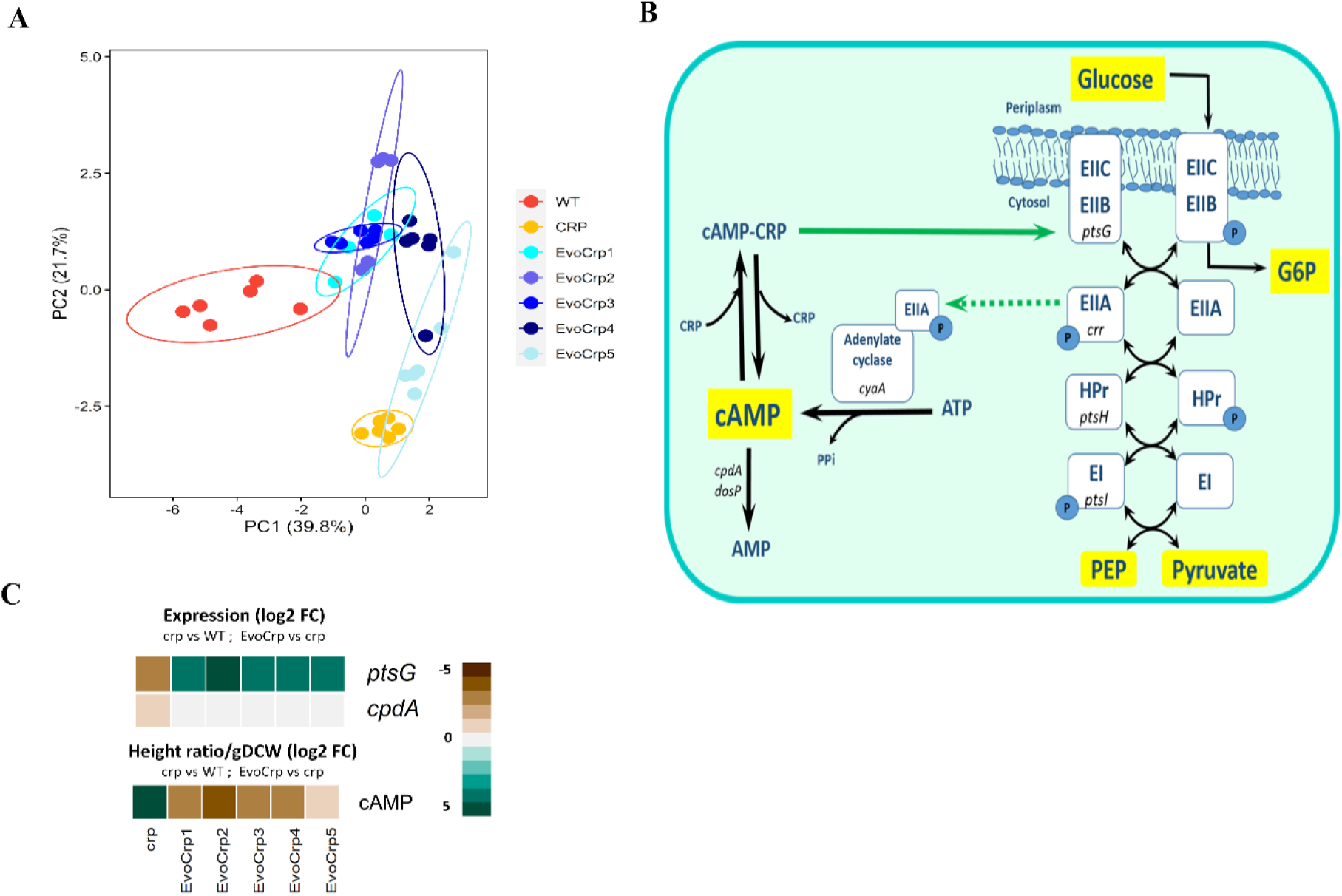
Fate of cAMP and global metabolome of the strains. (A) A multi-variate statistical analysis using PLS-DA for 24 significant metabolites found in our analysis. The six dots for each strain indicate the three biological and two technical replicates. The circles indicate 95% confidence intervals. (B) Network diagram of PEP-PTS system for glucose uptake with schematic representation of the regulation on cAMP by the enzymes of PEP-PTS. Key metabolites of the pathway are shown in yellow blocks. The green solid arrow indicates positive gene regulation from our study and the green dotted arrow indicates positive regulation of activity from literature evidence. (C) Expression profile of the DE genes of PEP-PTS with the metabolite levels of cAMP. Fold change values for gene expression and metabolite levels are obtained by comparing Δ*crp* vs WT and EvoCrp vs Δ*crp*. Expression values are obtained from average of two biological replicates expressed as log2 FC. Metabolite levels are obtained from average of three biological and two technical replicates expressed at height ratio/gDCW (log2 FC).

We integrated the metabolite levels with its cognate gene expression profiles to unravel the strain-specific adjustments at key nodes of metabolic pathways. Modulation of glucose uptake in *E. coli* can be inferred from the pool size of the physiological signal molecule cAMP, an inducer of CRP activity (fig. 4B). To decipher whether the absence of CRP alters the accumulation of cAMP, we measured the intracellular cAMP levels. We found that the deletion of CRP leads to a ~55-fold higher (adj-P < 10^−9^) accumulation of cAMP compared to the WT (fig. 4C). Counterintuitively, we observed no change in *cyaA* gene expression, that generates cAMP from ATP (fig. 4B), known to be activated by the phosphorylated EIIA (*crr*) component of the PTS system (fig. 4C) (Deutscher et al., 2006). Also, *cpdA* gene, responsible for degrading cAMP, had lower expression in Δ*crp*. Thus, the absence of sequestration by CRP and reduced degradation resulted in the extremely high intracellular accumulation of cAMP in Δ*crp*. After evolution, we observed a ~8-fold decrease in cAMP levels across all EvoCrp strains relative to Δ*crp* that indicated an evolutionary restoration of cAMP levels albeit inefficiently that might be suggestive of optimization of ATP usage.

We reinforce that glucose transport in exponential phase in *E. coli* K-12 MG1655 is mediated majorly via the *ptsG* gene (Deutscher et al., 2006; Postma et al., 1993), as evidenced by the mutation profiles. The gene expression pattern of *ptsG* (fig. 4C) in EvoCrp strains corroborates with the nature of mutations in the promoter of *ptsG* gene. Since glucose uptake in *E. coli* involves group translocation with the donation of phosphate from PEP (fig. 5A), we sought to examine PEP levels (Deutscher et al., 2006; Postma et al., 1993). We observed an increase in phosphoenolpyruvate (PEP) concentration (aFC ~ 8-fold) in Δ*crp* compared to the WT (fig. 5B), consistent with the reduced expression of*ptsG* gene (aFC ~ 2.5-fold). Recovery of PEP levels was observed across the majority of the Evo*Crp* strains (aFC ~ 6-fold, vs Δ*crp)*. Akin to PEP, 3-phosphoglycerate (3PG) was found to be higher in Δ*crp* (aFC ~ 3-fold, vs WT) and its level was reduced in EvoCrp strains (aFC ~ 2-fold, vs Δ*crp*). Since PEP and 3PG levels attune to *ptsG* expression, they represent reliable indicators of carbon limitation (Brauer et al., 2006), and their recovery promptly sheds light on the restoration of carbon balance. PEP being a precursor of aromatic amino acids, we observed a higher concentration of phenylalanine levels (aFC ~ 1.4) in Δ*crp*, concomitant with an increase in PEP levels (fig. 5B). Conversely, gene expression of aromatic amino acid biosynthesis genes namely *aroF* (aFC ~19)*, tyrA* (aFC ~13) and *trpABCD* (aFC ~3) showed significant downregulation in Δ*crp* mutant. Such antagonism highlighted the known negative feedback regulation of amino acid biosynthesis (Reznik et al., 2017). However, EvoCrp strains retained corresponding levels of phenylalanine compared to Δ*crp*, despite lowered levels of PEP and upregulation of *aroF* gene expression (aFC >7). Pyruvate, an α-keto acid and end product of glycolysis, is a precursor of the several branched-chain amino acids alanine, valine, leucine, and isoleucine (fig. 6A). We found a reduction in valine (aFC ~ 1.5-fold) and an increase in leucine levels (aFC ~ 2-fold) in Δ*crp* compared to WT. In contrast, we observed, reduction in valine (aFC ~ 2.3-fold, fig. 6D) and no significant changes in leucine in the EvoCrp strains compared to Δ*crp*. Although studies have suggested that higher valine inhibits branched chain amino acid synthesis and leucine biosynthesis involves sequestration of the toxic metabolite, acetyl CoA (Neidhardt et al., 1991), it is unclear as to why these strains choose to retain lower levels of valine and higher levels of leucine.

**Figure 5.**
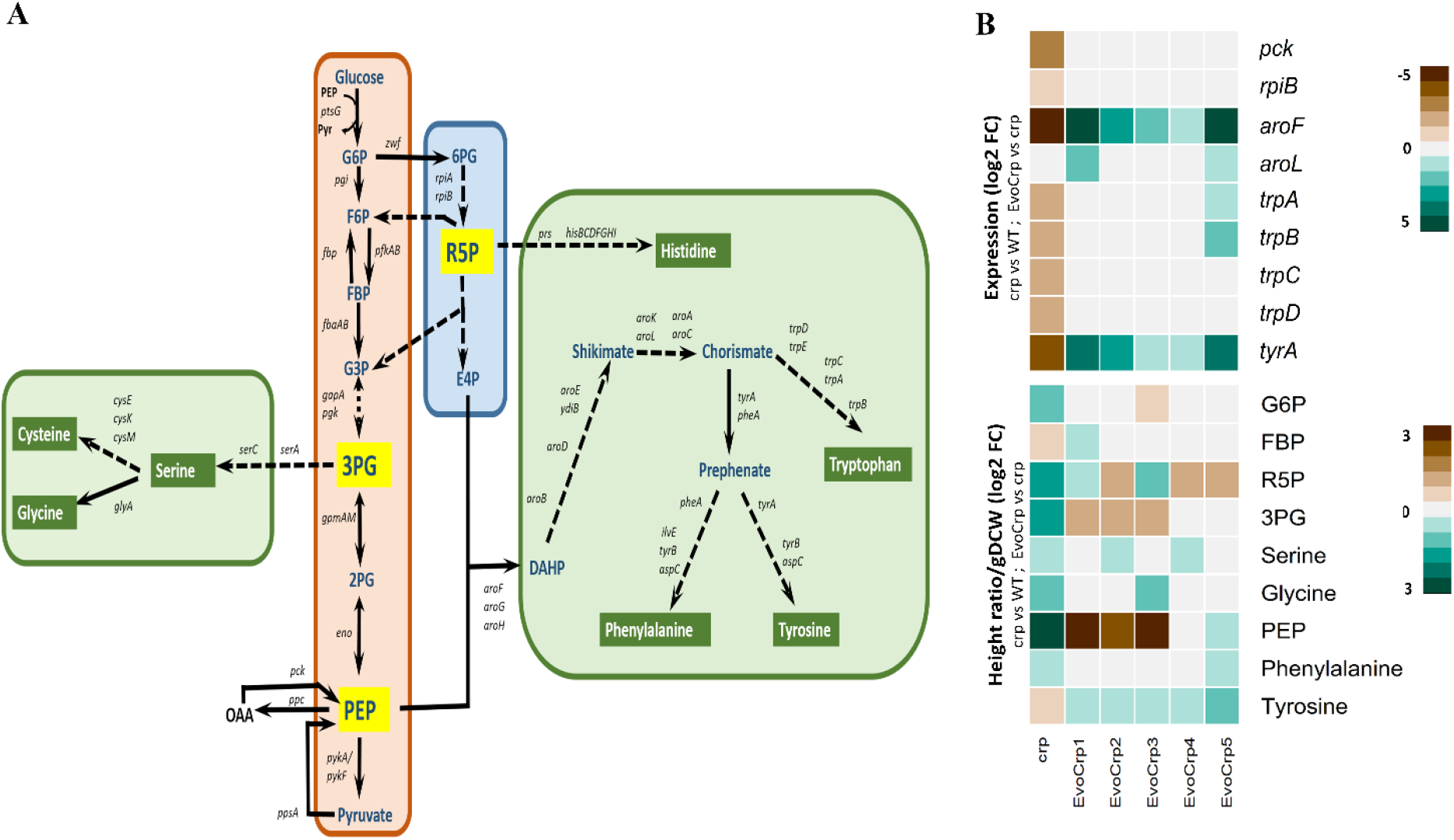
Integrated transcriptomics and metabolomics analysis at glycolytic and Pentose Phosphate Pathway (PPP) nodes. (A) A network diagram depicting glycolytic pathway, Pentose Phosphate Pathway (PPP) and amino acid biosynthetic pathways generating from the precursor metabolites of glycolytic and PPP pathway. (B) Expression profile of DE genes and metabolite levels significantly altered in the glycolytic and Pentose Phosphate Pathway (PPP) and amino acid biosynthetic pathways, were obtained by comparing Δ*crp* vs WT and EvoCrp vs Δ*crp*. Expression values are obtained from average of two biological replicates expressed as log2 FC. Metabolite levels are obtained from average of three biological and two technical replicates expressed as log2 FC of height ratio/gDCW.

**Figure 6.**
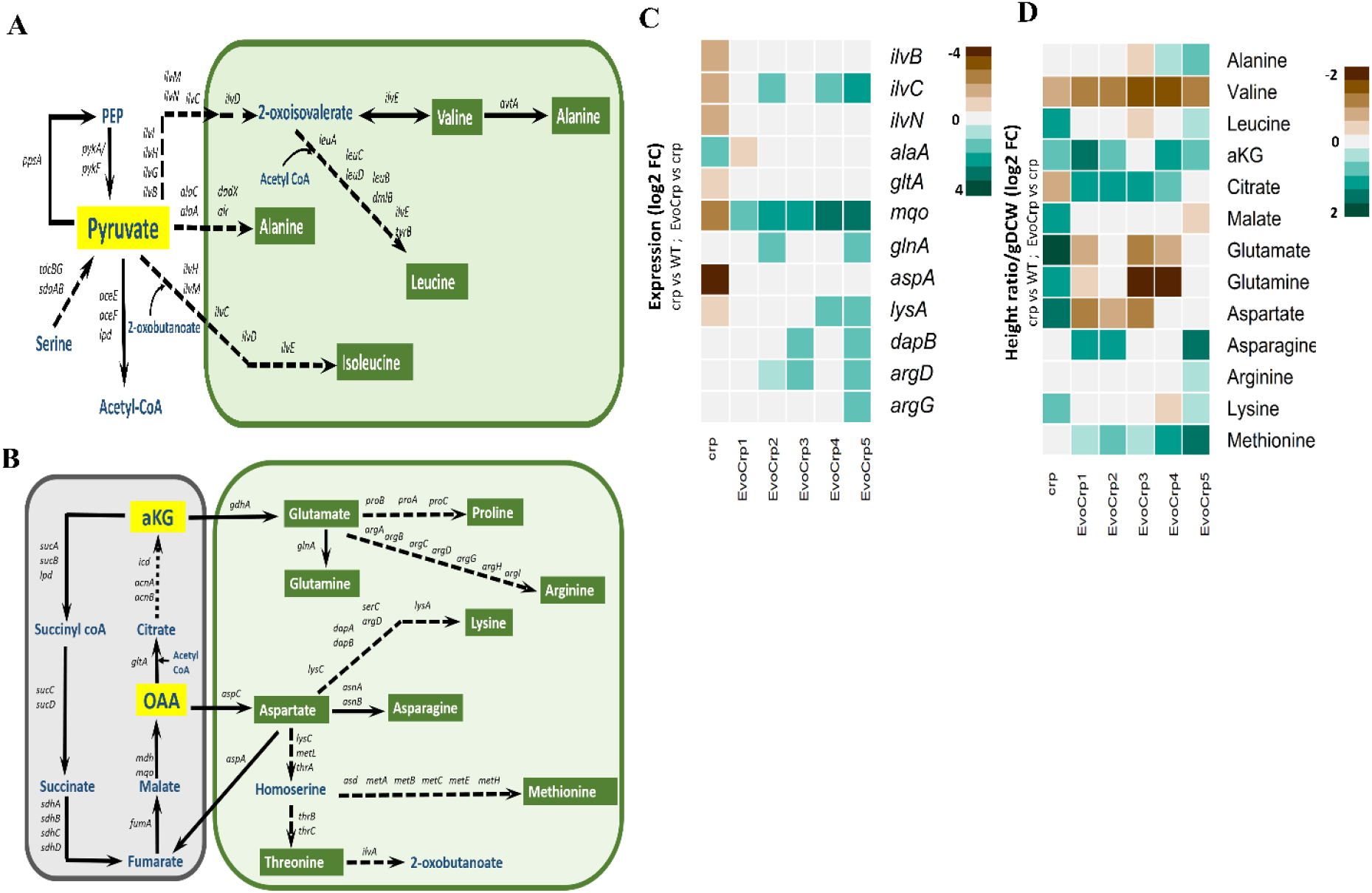
Integrated transcriptomics and metabolomics analysis at pyruvate and TCA cycle nodes. (A) A network diagram depicting branch points from pyruvate node, and amino acid biosynthetic pathways generating from the precursor metabolite pyruvate. (B) A network diagram depicting branch points from TCA cycle and amino acid biosynthetic pathways generating from the precursor metabolites of TCA cycle. (C) Expression profile of DE genes of pyruvate node, TCA cycle and amino acid biosynthetic pathways, were obtained by comparing Δ*crp* vs WT and EvoCrp vs Δ*crp*. Expression values are obtained from average of two biological replicates expressed as log2 FC. (D) Metabolite levels perturbed at the pyruvate node, TCA cycle and, acid biosynthetic pathways, were obtained by comparing Δ*crp* vs WT and EvoCrp vs Δ*crp*. Metabolite levels are obtained from average of three biological and two technical replicates expressed as log2 FC of height ratio/gDCW.

We also determined the concentrations of citrate and other α-keto acids like alpha-ketoglutarate (αKG) and oxaloacetate (OAA), which are key intermediates of the TCA cycle (fig. 6B). In Δ*crp*, citrate level was lower (aFC ~1.75) compared to WT in agreement with the reduced gene expression of *gltA* (fig. 6C-D), whereas its levels were restored after evolution despite no alteration in gene expression that might be attributed to increased TCA cycle activity in the EvoCrp strains (Eiteman et al., 2006). We observed a ~1.7-fold higher concentration of αKG in Δ*crp* compared to the WT as well as in EvoCrp compared to Δ*crp* (fig. 6D). The αKG accumulation has been known to indicate nitrogen limitation in the organism (Doucette et al., 2011). We speculate that the high intracellular levels of αKG in Δ*crp* as well as EvoCrp, therefore, indicated a possible scenario of nitrogen limitation prevalent due to loss of CRP and that evolution was unable to recover the limitation during the process of growth recovery. αKG is a precursor for the amino acids glutamate, glutamine, arginine, and proline. In Δ*crp* compared to WT, glutamate levels were upregulated (aFC ~3) whereas, in the EvoCrp strains compared to Δ*crp*, we observed reduced levels of glutamate (aFC ~1.6). OAA that can be interpreted from malate levels in the cell (Yuan et al., 2009), serves as a precursor of amino acids like aspartate, asparagine, lysine, threonine, and methionine levels (fig. 6B). Concomitant with the higher OAA or malate (aFC ~2-fold) levels, a ~2.5-fold higher aspartate and ~1.5-fold higher levels of lysine were seen in Δ*crp* mutant compared to WT. Despite no changes in OAA (malate) levels and ~1.75-fold reduction in aspartate concentration, we observed 1.6-fold higher asparagine and identical lysine levels in EvoCrp strains compared to Δ*crp*. Methionine, which showed no change in its concentration in Δ*crp* compared to the WT, was 1.6-fold high in all the EvoCrp strains compared to its parent Δ*crp*. Thus, concomitant with changes in precursors, we observed changes in proteinogenic amino acids, which elucidated an inefficient utilization towards its cellular objectives or protein biomass.

To summarize, we inferred that accumulation of key proteinogenic amino acids like phenylalanine, aspartate, glutamate, and lysine in Δ*crp* was ascribable to the impaired functioning of protein biosynthesis machinery observed in gene expression analysis. Further, in the EvoCrp strains, we observed increased accumulation of costly amino acids (considering only the number of activated phosphate bonds used in making the amino acid without the contribution of precursor synthesis itself)(Neidhardt et al., 1991) like methionine, asparagine, lysine, and arginine which were not restored to WT levels that seems to explain the restoration pattern seen in PLS-DA scores plot. Despite the restoration of protein biosynthesis machinery, the accumulation of these amino acids in EvoCrp was suggestive of the reallocation of ATP concordant with a reduction in unnecessary cAMP synthesis.

### Gene expression analysis of electron transport chain coupled to ATP synthesis

We assessed whether the reallocation of ATP was subjected to any changes in its synthesis genes. Hence, we investigated at the transcriptomic level any significant changes in the expression levels of the genes of the electron transport chain of *E. coli* that is responsible for the transfer of electrons to oxygen coupled to the generation of ATP (fig. 7A and 7C). We observed the downregulation of *cyoABCDE* genes (aFC~2) and the upregulation of *appABC* genes (aFC ~4.5) in Δ*crp* compared to the WT. The *cyoABCDE* genes are induced by high levels of oxygen and encode for the terminal oxidase cytochrome *bo*, the major route for oxygen channelling in aerobic respiratory conditions (Cotter and Gunsalus, 1992). On the contrary, *appABC* genes code for the terminal oxidase cytochrome *bd* and are expressed under low oxygenation levels (Atlungl and Brndsted, 1994). This reiterated the role of CRP in the proper utilization of oxygen coupled to changes in the TCA cycle (Nam et al., 2005; Zheng et al., 2004). In contrast, in EvoCrp compared to Δ*crp*, we observed restoration in these gene expression profiles. Additionally, the genes of ATP synthase complex, *atpABCDEFGH*, that are responsible for the synthesis of ATP (Neidhardt et al., 1991), were downregulated in the Δ*crp* compared to the WT. However, in EvoCrp strains relative to Δ*crp*, we observed an upregulation that indicated restoration of the ATP synthesis machinery.

**Figure 7.**
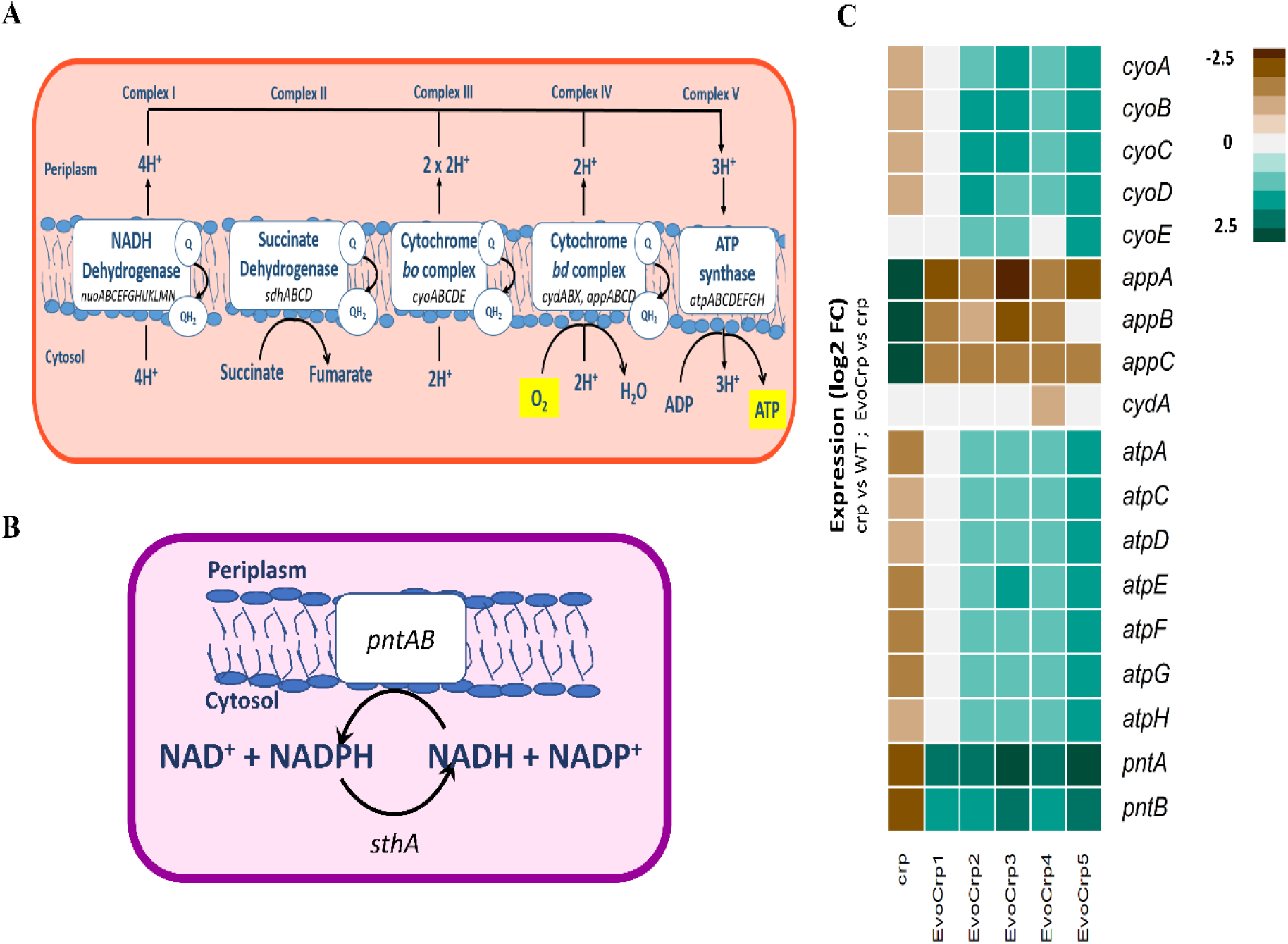
Gene expression data of oxidative phosphorylation and transhydrogenase genes. (A) A network diagram depicting the ETC and (B) the membrane bound transhydrogenases. (C) Expression profile of DE genes of ETC and the transhydrogenase genes, obtained by comparing Δ*crp* vs WT and EvoCrp vs Δ*crp*. Expression values are obtained from average of two biological replicates expressed as log2 FC.

Apart from ETC genes, we observed restoration in gene expression of membrane-bound transhydrogenase *pntAB* (fig. 7B and 7C), which was in agreement with the rapid growth recovery of EvoCrp strains. Previously, *pntAB* has been suggested to be necessary for anabolic NADPH formation whose expression profile levels have a strong correlation with the growth rate of the organism (Haverkorn van Rijsewijk et al., 2016; Sauer et al., 2004). To summarize, EvoCrp strains show restoration of oxidative phosphorylation and transhydrogenase expression profiles that are necessary for proper oxygen utilization and to balance the ATP and NADPH demand for anabolic processes to support enhanced growth.

### Model-based prediction of proteome allocation

An inherent property of *E. coli* is to tightly coordinate the metabolism and protein economy in the cell towards an optimal resource allocation favouring growth fitness (Erickson et al., 2017; Scott et al., 2010). In the light of observations from gene expression and metabolites, we sought to obtain insights into how necessary and unnecessary metabolic proteome are affected, which directly associates with the shifts in growth fitness. Towards this, we recalled a genome-scale ME-model (for Metabolism and Expression) that accounts for 80% of *E. coli* proteome (LaCroix et al., 2015; Lloyd et al., 2018; O’Brien et al., 2013; Utrilla et al., 2016), to assess how deletion of CRP affects the proteome allocation in the organism. The model was used to predict protein encoding genes that serves to explain the trade-offs in proteome allocation. The model predicted genes along with transcript per million (TPM) calculations were employed to depict the “utilized ME” (uME) and “non-utilized ME” (nonME) proteome fractions specific to aerobic glucose metabolism in *E. coli* K-12 MG1655 (supplementary file 1). We observed a reduction in uME fraction and an increase in nonME (glyoxylate shunt, osmotic stress-induced, and amino acid degradation genes) fraction in the Δ*crp* strain indicating a reduction in genes aligning with growth fitness and increase in genes related to hedging mechanisms respectively (supplementary fig. S17). Intuitively, our data revealed upregulation in uME fraction and downregulation in nonME fraction in EvoCrp strains to enhance their growth fitness. Also, a lower fraction of TCA cycle genes and secondary glucose transporters (*man* and *mal* genes) were found to be consistent across all the EvoCrp strains, to mitigate the unnecessary proteome cost towards their synthesis. Thus, as an adaptive strategy to recover growth, the EvoCrp strains redirected the protein economy towards necessary processes. Overall, changes in metabolic and unnecessary metabolic proteome towards biomass synthesis, the proteome allocation outline such trade-offs fundamental for balanced exponential growth (Scott et al., 2010).

### Physiological characterization agrees with the underlying molecular mechanism that defines growth

To evaluate how the changes in the transcriptome and the metabolome has directly impacted the phenotype of the organism, we characterized the growth rate, glucose uptake rate (GUR), acetate production rate (APR), oxygen uptake rate (OUR), and biomass yield (supplementary table S3). We observed a marked reduction in growth rate (~57%) in Δ*crp* strain compared to WT (fig. 8A). The GUR and OUR in Δ*crp* were both significantly reduced by ~56%, compared to the WT (P < 0.05, Student’s t-test) (fig. 8B and 8C), which can be attributed to the reduction in gene expression of *ptsG* and oxidative phosphorylation, respectively. Similar trends were also observed in APR as well, despite no changes in its gene expression (fig. 8D). Conversely, we observed increase in growth rate (112%, P < 0.05; fig. 8A) as well as GUR (~130%, P < 0.05; fig. 8B) and APR (~115%, P < 0.0 5; fig. 8D) in all EvoCrp strains compared to the Δ*crp*. This is in contrast to the previous study wherein, deletion of CRP in the ancestral strains did not cause any significant decrease in the glucose uptake rate and, *ptsG* gene gained CRP dependence only over time in their evolution experiment (Lamrabet et al., 2019). The OUR in EvoCrp showed variability in its increase compared to Δ*crp* ranging from ~99% to ~155% (fig. 8C). Further, we calculated the pairwise correlation of growth rate with GUR (Pearson correlation coefficient, r = 0.96, P < 10^−3^) and growth rate with OUR (Pearson correlation coefficient, r = 0.92). The strong correlation between them indicated that lowered GUR and lowered OUR might have resulted in an overall reduction of growth rate in the Δ*crp* strain (supplementary fig. S18A-S18B). In summary, our data suggested that all the parallel populations converged to phenotypes similar to WT at the end of ALE within ~100 generations.

**Figure 8.**
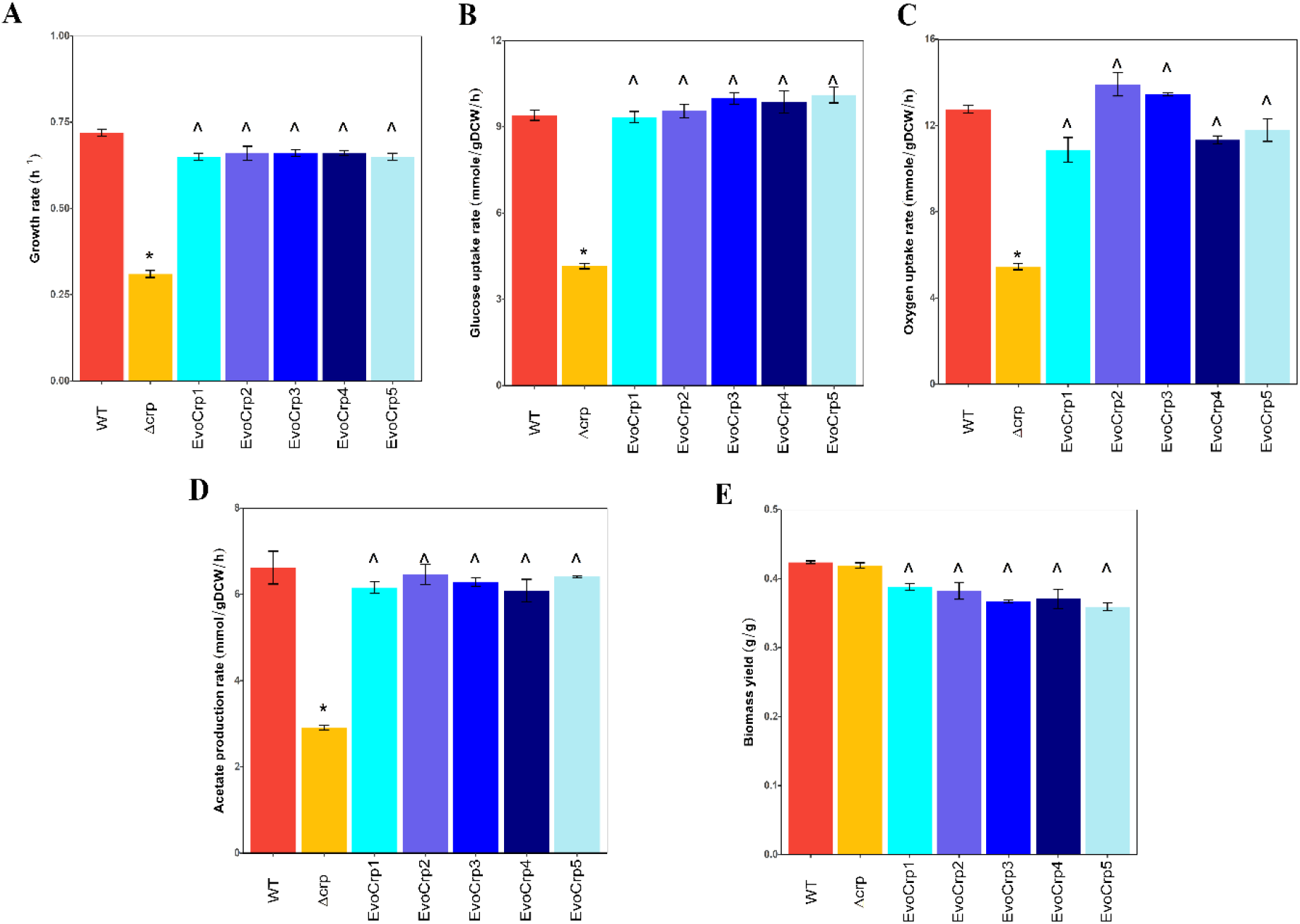
Phenomic studies of the strains. Bar-plot depicting the (A) growth rates, (B) glucose uptake rates (GUR), and (C) oxygen uptake rates (OUR), (D) acetate production rates (APR), and (E) biomass yields of the WT, Δ*crp* and the EvoCrp strains. The error bars represent an average of growth rates obtained from three biological replicates. The error bars indicate the standard error across the replicates. Significance of decrease in Δ*crp* vs WT is shown by asterisk whereas significance of increase in EvoCrp vs Δ*crp* is shown by caret (P < 0.05, Student’s t-test).

The biomass yield for each of the strains was determined by normalizing the growth rates with their specific glucose uptake rates. While the WT and Δ*crp* strains showed similar biomass yields (0.41-0.42 gDCW/gGlucose), there was a consistent decrease in the biomass yield (~0.37 gDCW/gGlucose) of EvoCrp strains that highlighted their inefficiency in optimally directing the carbon towards biomass (fig. 8E). Correlation of the measured biomass yield and growth rate (Pearson correlation coefficient, r = −0.87, P < 0.05, EvoCrp and Δ*crp*) showed the divergence of evolved strains away from Δ*crp* and WT (supplementary fig. S18C). Since changes in biomass yields can be a consequence of alterations in ATP maintenance (Pirt, 1982; Utrilla et al., 2016) (ATPM, i.e., ATP used for processes other than growth and protein synthesis), we predicted ATPM yields for each of the strains. The EvoCrp strains had a significantly higher ATPM compared to the WT as well as Δ*crp*, which was in agreement with the accumulation of costly amino acids (supplementary fig. S18D). This phenomenon of lowered efficiency of carbon substrate allocation towards growth reinforced the rate-yield trade-off mechanism prevalent in ALE-adapted strains (Cheng et al., 2019).

## Discussion

Adaptive mechanisms overarching genetic and metabolic regulatory networks are fundamental in conferring fitness advantages to strains when evolved in response to perturbations in its internal or external environments (Dragosits et al., 2013; Rodŕiguez-Verdugo et al., 2016; Sandberg et al., 2014; Utrilla et al., 2016; Zorraquino et al., 2017). In this study, we elucidated the pleiotropic changes due to the mutations in the *ptsG* promoter that enables the rapid growth of the strains when evolved in the absence of CRP. Specifically, we report restoration of precise functioning of protein biosynthesis machinery coupled with finely tuned proteome allocation and metabolic rewiring of ATP towards the synthesis of costly amino acids away from the wasteful cAMP synthesis in addition to enhanced rates of glucose uptake and related physiological traits, that overall results in growth fitness.

During evolution, mutations occurred predominantly in the *ptsG* promoter that generated additional “Pribnow-box” like sequences to enhance the affinity of RNA polymerase sigma 70 towards *ptsG* gene. Indeed, the lack of interplay of other regulators seen from *in vivo* binding studies and RNA polymerase sigma factor reallocation seen from transcriptome analysis supports this basis. Notably, the mutations resulted in increased *ptsG* gene expression and thereby glucose uptake, emphasizing the role of *ptsG* promoter mutations in resolving the bottleneck caused by loss of CRP. Such a phenomenon was also observed in a previous study, thereby implying genetic parallelism across different *E. coli* sub-strains (Lamrabet et al., 2019). Despite such similarities, the difference in regulatory control of CRP on *ptsG* gene and the profound differences observed in genomic and phenomic states concurs with the genetic background of the parent strains. The secondary glucose transporters, known to function in limiting glucose or pH altered conditions (Steinsiek and Bettenbrock, 2012), did not harbour any mutations *per se*, which might be attributed to serial passaging in the mid-exponential phase and constant pH during ALE. Thus, we corroborate with previous studies (Deutscher et al., 2006; Postma et al., 1993) that *ptsG* is the sole major transporter of glucose in *E. coli* K-12 MG1655 during its exponential growth.

We also elucidate here the molecular events underlying impaired growth caused by the deletion of CRP that delineates the coordination between the global regulator and several growth-rate dependent factors (RNA polymerase, ribosome, and metabolites) (Berthoumieux et al., 2013; Klumpp et al., 2009). We report that the deletion of CRP caused a significant decrease in the glucose and oxygen uptake rates, which correlated strongly with the decreased growth rate. We additionally emphasized the indirect effects of CRP deletion on protein biosynthesis machinery, as evidenced by their metabolic and ribosomal gene expression profiles as well as the accumulation of proteinogenic amino acids that were not being utilized for protein biomass. Recent studies have reported tight coordination of ribosomal protein expression with the growth rate of the organism (Scott et al., 2014, 2010). Biosynthesis of ribosomes required for protein translation involves huge proteome investment (Scott et al., 2010; Zhu and Dai, 2019). For slow-growing cells in stressed environmental conditions, the organism produces more ribosomes to hedge for unfavourable conditions (Zhu and Dai, 2019). This, in turn, limits the ribosomes available for synthesis for metabolic proteins manifested as an increase in unnecessary ribosomal proteins and a reduction in translational efficiency thereof (Dai et al., 2018).

Concurrently, our predicted system-wide proteome fraction of Δ*crp* mutant demonstrated a wasteful allocation of proteome towards stress or hedging functions. Evolved strains rebalanced the proteome by increasing the growth-promoting proteome fraction and by mitigating the proteome cost associated with the synthesis of expensive proteins, such as enzymes of the TCA cycle. However, to meet the energetic demands of the faster-growing strains, we suggest that the TCA cycle enzymes might have gained a higher *in-vivo* maximal rate observed from higher levels of TCA cycle intermediates, a phenomenon predicted to be occurring in adaptive evolution (O’Brien et al., 2016). In addition to lowered proteome fractions of secondary glucose transporters in the evolved strains, our findings emphasized the fine-tuning of the proteome allocation in response to its metabolic demands.

It is well known that growing cells maintain an optimal cAMP level necessary for proper carbon sensing (You et al., 2013) and hence ATP optimization. We observed a high accumulation of cAMP in the absence of sequestration by CRP. On the contrary, evolved strains show restoration of their cAMP level though inefficiently, thereby partially alleviating ATP wastage. We speculate that this surplus ATP pool conserved from reduced cAMP synthesis was invested towards the synthesis of costly proteinogenic amino acids that were found to accumulate in the evolved strains. This might be partially responsible for the higher ATPM yield resulting in an overall reduction in the biomass yields observed in the evolved strains. It should be noted that such accumulation in evolved strains can also be ascribable to protein degradation which is energetically expensive (Jozefczuk et al. 2010; Barupal et al. 2013). Nevertheless, such inefficiencies involving subpar utilization of carbon towards biomass in the evolved populations limit their ability to grow as energetically optimal as the wild-type. Such inefficiencies involving subpar utilization of carbon towards biomass in the evolved populations limit their ability to grow as energetically optimal as the wild-type. Overall, we observed that Δ*crp* efficiently allocated carbon but not proteome towards biomass synthesis, whereas, the evolved strains were able to efficiently allocate only the proteome towards biomass synthesis.

In summary, we have elucidated in detail the system-wide pleiotropic effects manifested due to the mutations in the promoter of *ptsG* gene. The current ALE study using multi-omics has revealed mechanistic insights into the inherent systemic constraints that facilitate the final phenotypic response in conjunction with the selected mutations in the evolved strains to overcome the defects due to loss of global regulator. We observed that growth fitness in the adaptively evolved strains in comparison to its parent Δ*crp* is enhanced due to modulation of the metabolic and ribosomal gene expression along with alterations of several central carbon metabolites namely PEP, α-ketoacids, cAMP, and proteinogenic amino acids that facilitate precise coordination of protein biosynthesis machinery with metabolism. Further, we revealed that the evolved strains possessed finely tuned proteome allocation that favour growth over uneconomical hedging strategies and mitigation of costly proteome fractions. Our findings based on proteome allocation in parent and evolved strains bear striking similarities with proteome sector changes, that represent a corollary to resource allocation defined by growth laws (Scott et al., 2014, 2010). Future studies characterizing the genotype-to-phenotype relationship using a multi-omics approach would expedite our understanding of microbial evolution across diverse conditions.

## Materials and Methods

### Strains

An *E. coli* K-12 MG1655 (CGSC#6300), was used as the parent strain in this study and all the other strains used in this study were derived from this strain (supplementary table S1). We constructed Δ*crp* knockout in this genetic background by λ-Red mediated recombination (Datsenko and Wanner, 2000), using plasmids pKD46, pKD13, and pCP20. After strain verification, glycerol stocks were made and stored at −80°C.

### Physiological characterization in a bioreactor

For transcriptome, metabolome and phenotype characterizations, cells were grown in 500 mL bioreactor (Applikon) (planktonic state, batch culture) containing 200 mL M9 media (6 g/L anhydrous Na_2_HPO_4_, 3 g/L KH_2_PO_4_, 1g/L NH_4_Cl, 0.5 g/L NaCl + 2 mM MgSO_4_ + 0.1 mM CaCl_2_), with 2g/L glucose and 40 mM MOPS. Briefly, cells from glycerol stocks were plated out on LB agar plate and a single colony was inoculated in LB media. A fixed volume of 100 μL cells was used to inoculate 50 mL preculture M9 +40 mM MOPS media with 4 g/L glucose which was grown overnight in a shake flask at 200 rpm in a 37°C (Eppendorf) incubator. The preculture cells, while still in exponential phase, were centrifuged and washed with M9, and inoculated in a bioreactor containing 200 mL M9 media with 2g/L glucose such that the start OD of all the cultures were ~ 0.05 OD. The temperature of the bioreactor was maintained at 37°C and the pH of the media was maintained at pH 7.2 using 40 mM MOPS buffer to prevent the effects of pH change on growth. Aeration was done by sparging air in the bioreactor at 700 mL/min and at all times the dissolved oxygen (DO) levels were maintained above 40% saturation using mass flow controllers (MFC). Growth was monitored by collecting samples at regular intervals and measuring the optical density (OD) at 600 nm in a spectrophotometer (Thermo Multiskan GO) until the organism reached its stationary phase. The growth rate was calculated from the slope of the linear regression line fit to ln(OD600) vs. time (in hrs) plot in the exponential phase. The dry cell weight (DCW) was determined from the O.D.600 values by using the experimentally derived relationship that 1.0 O.D.600 corresponds to 0.44 gDCW*h^−1^. The phenotypic characterizations were done for three biological replicates (n = 3) across the exponential phase. The transcriptome and metabolomics characterizations were performed in the mid-exponential phase (0.6-0.7 OD). Samples were also collected during regular intervals to determine the rate of glucose uptake as well as the rate of secretion of extracellular metabolites. The samples were centrifuged and the supernatant was collected which was then used to determine the concentrations and rates using HPLC (Agilent 1200 Series) equipped with Bio-Rad Aminex HPX-87H ion exclusion column with 5mM H2SO4 as the mobile phase. The column was maintained at a temperature of 50°C and the flow rate at 0.6 mL/min. The specific rates were calculated from the change in substrate concentration over time, normalized to the biomass of each strain (Sauer et al., 1999). The biomass yield for each of the strains was determined by normalizing with their specific glucose uptake rate (gDCW/gGlucose). Oxygen uptake rate was measured using a dissolved oxygen probe (Applikon) while growing the cultures in the bioreactor during its exponential phase. All physiological measurements were checked for statistical significance using unpaired two-tailed Student’s t-test.

### Adaptive Laboratory Evolution Protocol

Adaptive evolution of replicate populations of Δ*crp* was carried out in shake flasks (planktonic state, batch culture) with M9 minimal media with 2g/L glucose and 40 mM MOPS at 37°C. MOPS was added to maintain the populations at a constant pH during evolution (supplementary fig. S2). All the replicate cultures were passaged serially into fresh media strictly in the mid-exponential phase to ensure that fitness gains occur primarily via increased growth rates. This also prevents the cultures from entering the stationary phase where glucose-limited and thereby avoids complexities associated with the onset of the stationary phase. As the growth rate of the organism changed during evolution, the volume of culture passaged was adjusted to prevent entry into the stationary phase. Glycerol stocks were made during each passage and PCR verified for any cross-contamination. This procedure was followed for 10 days (approximately 100 generations) and after it had reached a stable growth and no further increase in growth rate was observed, ALE was terminated. Genotype and phenotype characterizations of the evolved replicates were done for the population and not for individual clonal samples to ensure that all the traits of the population are taken into consideration. Characterization at the population level is a more efficient and feasible approach to understand the underlying mechanisms of evolution in an unbiased manner as it better reflects the properties of the population as a whole (McCloskey et al., 2018b).

### Whole Genome Resequencing (WGS)

Genomic DNA from all the endpoint populations was extracted using GenElute Bacterial Genomic DNA Kit (NA2120; Sigma-Aldrich, St. Louis, MO) using the manufacturer’s protocol. The integrity of the extracted genomic DNA was analyzed by running it on an agarose gel and the quality was assessed and quantified using Multiskan GO (Thermo Scientific). The genomic DNA library was prepared using Illumina TruSeq DNA Nano Kit. The quality of the libraries was checked using Agilent Bioanalyzer. The libraries were then sequenced from both ends (paired-end) on Illumina HiSeq250 platform with 2 × 100 cycles. All the samples had an average of 275X mapped coverage.

The raw reads obtained from the sequencer were trimmed using CUTADAPT to remove TruSeq adapter sequences. The breseq pipeline (Deatherage and Barrick, 2014) was used to identify the point mutations (SNPs), insertions, and deletions mapped to *E. coli* genome K-12 MG1655 (GenBank accession no. NC_000913.3). Breseq was run using the -p option (for population samples) with default parameters to identify mutations present in only a subset of the population at a frequency of less than 100%. Mutations that had 100% frequency were assumed to be parental mutations and were not considered. Only the mutations predicted with high confidence under the category “predicted mutations” were further analyzed in this study.

### Mutation Validation

To determine the causality of the mutations detected by WGS, the mutations were introduced into the Δ*crp* background using *in vivo* site-directed mutagenesis (Kim et al., 2014). However, to limit the scale of this study and given the consistent occurrence of mutations in the Fis binding region, only the IG116 mutation (SNPs) detected in the promoter of *ptsG* gene in the EvoCrp1 strain was selected for validation. Additionally, IG116 mutation was also investigated for Mlc binding. To construct the strains, scarless editing of the *E. coli* K-12 MG1655 Δ*crp* genome was done using a two-step recombination method using pSLTS plasmid (Addgene plasmid #59386). The primers used in this method are given in Supplementary Table S2. After the construction of the strains, they were sequenced to verify the correct introduction of the SNPs in the genome. This strain was further used to add 3X FLAG-tag to Fis protein or Mlc protein as described in the section below.

### ChIP qPCR

A 3X FLAG-tag was added to the C-terminus of the Fis or Mlc protein using pSUB11 plasmid (Uzzau et al., 2001) as the amplification template. The primers used for amplification for tagging are given in Supplementary Table S2. The amplified construct was then introduced into *E. coli* K-12 MG1655 WT, Δ*crp* and IG116-Δ*crp* strain. The constructed strains were verified by PCR and Sanger sequencing. ChIP experiment was carried out using the protocol as described previously (Kahramanoglou et al., 2011). The DNA samples immune-precipitated by this method were recovered by PCR purification and were quantified by qPCR using primers for the specific as well as non-specific region (*frr*). Fold change in IP-ed DNA compared to mock DNA was calculated as 2^−ΔΔCt^ as described previously (Mukhopadhyay et al., 2008).

### RNA extraction and enrichment of mRNA

For each sample/strain, RNA extraction of two biological replicates was performed. The cells were grown till the mid-exponential phase and then 50 mL cells were harvested by centrifugation. RNA extraction was done using the TRIzol-chloroform method as previously described (Rio et al., 2010; Srinivasan et al., 2015). DNase treatment was done to remove any DNA contamination after which enrichment for mRNA was done using the MicrobExpress Kit following the manufacturer’s protocol. Single-end, strand-specific libraries for RNA sequencing were prepared using NEBNext Ultra II Directional RNA library kit. The quantity of the mRNA was assessed using Multiscan GO (Thermo Scientific) Nanodrop and the quality, as well as integrity, was checked using BioAnalyzer. The sequencing was carried out on HiSeq 2500 Rapid Run Mode using a 1×50 bp format.

### Transcriptome data analysis

The raw files were first trimmed using CUTADAPT to remove adapter sequences and low-quality reads. The reads were then mapped to the *E. coli* genome K-12 MG1655 (GenBank accession no. NC_000913.3) using BWA (Li and Durbin, 2009). The sam file generated was then converted to compressed bam format using Samtools (Li et al., 2009). FeatureCounts (Liao et al., 2014) was used to assign counts at the gene level using the reference genome provided in the GTF format. Ecocyc database (Keseler et al., 2017) (version 21.5) was used to retrieve the annotations for 4662 genes. Differential gene expression (DGE) was analyzed from the raw counts by EdgeR (Robinson et al., 2009) after removing genes having less than 10 reads. The genes that showed ≥ 2-fold change in expression (in both directions) and had adjusted P < 0.05 (Benjamini-Hochberg) were considered for further analysis. The DEGs were enriched for metabolic pathways using KEGG pathway classification (Minoru Kanehisa and Susumu Goto, 2016) as defined in Proteomaps (www.proteomaps.net). The mapped genes were represented as Voronoi treemaps (version 2.0) (Liebermeister et al., 2014). The significance of the upregulated and downregulated genes within each category was validated using a hypergeometric test with P-value correction using Benjamini Hochberg in R (R Core Team 2019). For each up/down regulated category, we arbitrarily choose atleast five DE genes enriched to be considered for significance analysis. Those DE genes which do not have an assigned “Accession ID” (Ecocyc Version 21.5) such as phantom genes were excluded from the above analysis.

For sigma factor enrichment analysis, the DE genes enriched by KEGG pathway classification were further enriched based on regulators using data available in EcoCyc and Regulon DB (Santos-Zavaleta et al., 2019). The upregulated and downregulated genes belonging to each regulator were then validated using a hypergeometric test in R (P-value < 0.05). Only those sigma factors which regulated atleast 10% of the KEGG enriched upregulated and downregulated DE genes, were retained for this over-representation analysis. Pearson correlation of log2 fold changes of EvoCrp vs WT and Δ*crp* vs WT and statistical significance were performed in R (R Core Team 2019).

### Metabolomics

#### 1. Chemicals for metabolomics

LC-MS grade methanol, acetonitrile, and ammonium hydroxide (≥ 25% in water) were purchased from Honeywell. Analytical grade chloroform, HPLC-grade water, and LC-MS grade ammonium acetate were purchased from Sigma. All metabolites used as external standards were purchased from Sigma. Uniformly labelled U-^13^C (> 99% purity) glucose was purchased from Cambridge Isotope Laboratories.

#### 2. Extraction of metabolites

Metabolite samples were harvested in the mid-exponential phase in biological triplicates and technical duplicates. A Fast-Cooling method (Fu et al., 2015; Hollinshead et al., 2016) was used to quench the harvested cells as reported previously. Briefly, ~10 mL culture (~ 6-7 OD cells) was directly poured into 2 mL chilled (4°C) M9 (without glucose) in a 50 mL falcon tube. The tube was then dipped in liquid nitrogen for 10 secs to bring down the sample temperature to 0°C. To prevent the formation of ice crystals, the sample was vigorously agitated with the help of a digital thermometer. Samples were then immediately centrifuged at 0°C, 7800 rpm for 5 min. The supernatant was discarded and the pellet was snap-frozen in liquid nitrogen and stored at −80°C until metabolite extraction was done.

For extraction of metabolites, the sample pellet was dissolved in 700 μL chilled (−80°C) methanol: 300 μL chilled (−20°C) chloroform (Hollinshead et al., 2016), followed by snap freezing in liquid nitrogen and homogenization using a hand-held pestle (Sigma #Z359971) all within 2 min. ^13^C-labelled extracts as internal standards were generated for quantification of key metabolite pool sizes using an isotope-based dilution method (Wu et al., 2005). The samples were spiked at the earliest stage of extraction with a fixed volume of internal standard taken from the same batch. The sample tubes were then placed overnight on a Thermo-shaker maintained at 0°C and 400 rpm. To the sample tubes, 500 μL of 2% ammonium hydroxide (4°C) prepared in HPLC-grade water was added and incubated on ice for 10 min to enable phase separation. The tubes were then centrifuged at 4°C, 13000 rpm for 15 min and the aqueous layer was collected in a chilled 1.5 mL tube. The samples were then completely dried in a vacuum concentrator and stored at −80°C. Before analysis in LC-MS/MS, samples were reconstituted in 100 μL chilled (−20°C) acetonitrile: buffered water (60:40, v/v), centrifuged at 4°C for 10 min, and the supernatant was transferred to pre-chilled glass vials. Buffered water consisted of 10 mM ammonium acetate, pH 9.23 adjusted with ammonium hydroxide prepared in HPLC-grade water (Teleki et al., 2015). The volume of the pooled internal standard was standardized such that the external standard and internal standard peak height differed less than 5-fold (Bennett et al., 2009). Metabolites were extracted from three biological and two technical replicates (n = 6).

#### 3. LC-MS/MS settings

The samples were analyzed using a high-resolution mass spectrometer in Orbitrap Q Exactive Plus (Thermo) equipped with a SeQuant ZIC-pHILIC column (150 mm x 2.1 mm x 5-micron packing, Merck) and a ZIC-pHILIC guard column (20 mm x 2.1 mm x 5-micron packing, Merck) under alkaline mobile phase conditions with ESI ion source. The ESI was operated in positive (M+H)^+^ and negative (M-H)^−^ polarity switching mode. The spray voltage was set at 4.2 kV and 3.5 kV for the positive and negative modes respectively. The temperature was maintained at 300°C and 320°C for the ion transfer capillary and probe heater respectively. A heated electrospray ionization probe II (H-ESI-II) probe was used with the following tune parameters: sheath gas, 29; auxiliary gas, 7; sweep gas, 0; S-lens at 45 arbitrary units. A full scan range of 66.7 to 1000 m/z was applied for positive as well as negative modes and the spectrum data type was set to profile mode. The automatic gain control (AGC) target was set at 1e6 with a resolution of 70,000 at 200 m/z. Before analysis, cleaning of the LC-MS system and ion transfer capillary (ITC) along with mass calibration was done for both positive and negative ESI polarities by using Thermo Calmix solution along with MS contaminants to take into account lower mass ranges. The signals of compounds 83.06037 m/z (2 x ACN+H) and 119.03498 m/z (2 x Acetate-H) were selected as lock masses for positive and negative mode respectively with lock mass tolerance of 5 ppm (Zhang et al., 2014).

The mobile phase for chromatographic separation comprised of non-polar phase A (acetonitrile: water mixed in the ratio 9:1, 10 mM ammonium acetate, pH 9.23 using ammonium hydroxide) and polar phase B (acetonitrile: water mixed in the ratio 1:9, 10 mM ammonium acetate, pH 9.23 ammonium hydroxide). A linear gradient with flow rate of 200 μL/min was set as follows: 0-1 min: 0% B, 1-32 min: 77.5% B, 32-36 min: 77.5% B to 100% B, 36-40 min: hold at 100% B, 40-50 min: 100% B to 0% B, 50-65 min: re-equilibration with 0% B (Teleki et al., 2015). An injection volume of 5 μL was used for all the samples and standards.

#### 4. Metabolomics data analysis

The data from the machine was processed using the software package Xcalibur 4.3 (Thermo Fisher Scientific) Quan Browser. A semi-quantitative analysis was performed using peak heights of precursor ions with signal/noise (S/N) ratio more than 3, a retention time window of less than 60 secs, and less than 5 ppm mass error. The peak heights of the samples were normalized to the peak height of the internal standards to obtain a height ratio. Only those metabolites were retained for analysis, which had naturally occurring ^12^C peak height less than ~10% of the ^13^C-labelled peak height in the internal standard. MetaboAnalyst (Xia et al., 2009) was used for identifying statistically significant metabolites and for Partial Least Square Discriminant Analysis (PLS-DA) on biomass normalized and g-log transformed metabolite concentrations. Missing value imputation was performed using SVD impute function. The concentration of metabolites is expressed as height ratio normalized to biomass (as height ratio/gDCW). Metabolite levels with false discovery rate (FDR) < 0.05 and ≥ 1.2-fold change in concentrations (in both directions) were considered for further analysis. For PLS-DA, out of the 40 metabolites, only those that were statistically significant in all the three conditions, i.e., Δ*crp* vs WT, EvoCrp vs Δ*crp*, and EvoCrp vs WT, were considered for analysis. In the heatmaps generated based on log2 fold change of metabolites, only those metabolites with FDR < 0.05 in atleast one condition were retained for comparative analysis (the same metabolite with FDR > 0.1 in other conditions were not considered).

### ME-model simulations

The ME-model (Lloyd et al., 2018; O’Brien et al., 2013) was simulated to generate a gene list for analysis, by constraining the glucose uptake rates in a range as reported previously (LaCroix et al., 2015). Any gene predicted to be expressed in any of the 20 simulations was classified as “utilized ME”; genes within the scope of the ME-model but not expressed or having very low expression (low protein synthesis flux) are classified as “non-utilized ME”; Next, using the differentially expressed genes in atleast one condition (i.e., Δ*crp* vs WT, EvoCrp vs Δ*crp* and EvoCrp vs WT) were combined and enriched for utilized ME or non-utilized ME gene list followed by summing the transcriptome fractions of all utilized ME and non-utilized ME genes respectively. The transcriptome fraction was determined from the product of TPM (transcript per million) and gene length divided by the sum-product of these calculated over all the genes (Utrilla et al., 2016). TPM (Wagner et al., 2012) for all the 4662 genes were calculated using the raw counts (genes < 10 reads were not considered). As the proteome fractions were not measured experimentally, we assumed that the increase in these protein-coding transcriptome fraction corresponds to an increase in proteome fractions. Hence, utilized and non-utilized ME genes are metabolic genes corresponding to utilized and non-utilized metabolic proteome, respectively (supplementary file 1).

### Calculation of non-growth ATP maintenance (ATPM)

The experimentally measured growth rates, glucose uptake, and acetate secretion rates were used as flux constraints in the iJO1366 metabolic model (Orth et al., 2011). Constrained-based flux balance analysis was performed using COBRApy (Ebrahim et al., 2013) by maximization of ATP maintenance (ATPM) reaction flux. This predicted ATPM flux was normalized to glucose uptake flux to calculate the yield (gATP/gGlucose) for each of the strains.

## Acknowledgments

We thank Jeffrey E. Barrick (Associate Professor, Department of Molecular Biosciences, The University of Texas at Austin) for his inputs in WGS and site-directed mutagenesis. We thank Edward Baidoo (Biochemist Research Scientist, Lawrence Berkley National Laboratory) for his valuable suggestions in metabolomics. We thank Awadhesh Pandit (Next Generation Genomics Facility, C-CAMP, NCBS) for providing genomics and transcriptomics services. We thank Dr. Mayuri Gandhi (Research Scientist, SAIF, IIT Bombay) for providing the LC-MS facility. We thank Prof. Pradeepkumar P. I. (IIT Bombay) for providing RT-PCR facility and Prof. Sarika Mehra (IIT Bombay) for providing vacuum concentrator for our use. We thank Colton Lloyd (Systems Biology Research Group, UCSD) for his inputs in ME-model simulations. We thank Reshma T. V. (NCBS) and Guhan K.A. (IIT Bombay) for their valuable technical inputs in ChIP-qPCR experiments. We also thank Dr. Pramod Rajaram S for his comments on the manuscript.

This work was supported by DST (Department of Science and Technology) fellowship/grant (SB/S3/CE/080/2015) and DBT (Department of Biotechnology) fellowship/grant (BT/PF13713/BBE/117/83/2015) awarded to K.V.V., DBT/Wellcome Trust India Alliance Intermediate Fellowship/grant (IA/I/16/2/502711) awarded to A.S.N.S. and DST-WOSA grant (SR/WOS-A/ET-58/2017) awarded to S.S.

A.P. acknowledges the Department of Science and Technology INSPIRE, Govt. of India for her fellowship (IF140914). M.S.I acknowledge the Department of Biotechnology, Govt. of India for his fellowship (DBT/IIT-P/323).

## Author Contributions

A.P. and K.V.V. designed the research; A.P. and M.S.I. performed research; A.P. and M.S.I. analyzed data; A.P., M.S.I., and S.S. performed computational modelling; A.P., M.S.I., and S.S. contributed to new reagents/analytical tools; K.V.V provided inputs on aspects of metabolism; A.S.N provided inputs on genome and transcriptome studies; A.P. and M.S.I. wrote the paper, A.S.N. and K.V.V. revised the paper. All authors read and agreed on the manuscript.

## Data availability

The whole-genome resequencing data presented this study are available at NCBI BioProject (https://www.ncbi.nlm.nih.gov/bioproject/) under accession no. PRJNA637503. The RNA sequencing data and the processed files from this study are available at NCBI Geo (https://www.ncbi.nlm.nih.gov/geo/query/acc.cgi?acc) under accession no. GSE152214. The metabolomics data presented in this study is available at the NIH Common Fund’s National Metabolomics Data Repository (NMDR) website, the Metabolomics Workbench, (https://www.metabolomicsworkbench.org) where it has been assigned Project ID (PR000957). The data can be accessed directly via it’s Project DOI: (10.21228/M80D7P).

