## Supplementary material for "Global pleiotropic effects in adaptively evolved *Escherichia coli* lacking CRP reveal molecular mechanisms that define growth physiology"

### Supplementary text:

Whole genome sequencing revealed a total of 11 unique mutations across all the EvoCrp strains. Most of these were single nucleotide polymorphisms (SNPs) with one or more of these mutations being present in more than one evolved strain. The mutation profile of no two evolved replicate populations were exactly the same. This indicates that each of them traversed through different evolutionary landscapes to arrive at a converging phenotypic response. Out of the 11 unique mutations, 8 were in the promoter region of the *ptsG* or in the coding region. Most of these mutations were SNPs (single nucleotide polymorphisms) and one 14bp duplication. A detailed analysis of each of the mutations show that these SNPs as well as the duplication resulted in additional "Pribnow-box" like sequences, thereby enhancing the binding affinity of RNA Polymerase for the promoter region, that in turn increases *ptsG* gene expression, as evident for the RNA-seq analysis of the EvoCrp strains. Mutation of the promoter region IG276, a C →T transition mutation (T**C**GTAA→ T**T**GTAA) 276 bp upstream of the start of *ptsG* gene, was the most prominent one being present in 4 out of 5 of the evolved replicates. Another prominent mutation in the promoter region is IG106 mutation that is 3 bp upstream of the start of *ptsG* gene. It is present in 2 out of 5 replicates and causes a G → A transition mutation (T**G**TAAT → T**A**TAAT). There are two other SNPs, IG116 (TTAT**T**T → TTAT**G**T) and IG250 (TAAA**G**T → TAAA**A**T) in the promoter region. Moreover, one of the evolved strains had the mutation IG98, a 14 bp duplication ( $[TCTGTGTAATAAAT]_2$ ) at 98 bp upstream of the *ptsG* start site. Also, EvoCrp2 showed a mutation upstream of the *ptsG* gene in the binding site of SgrS, a small regulatory RNA. SgrS inhibits *ptsG* post-transcriptionally by binding to *ptsG* mRNA, causing translational silencing and indicates a negative regulation of sgrS on *ptsG* gene. We hypothesize that the mutation at SgrS binding site might prevent the binding of SgrS to *ptsG* mRNA, thereby preventing transcriptional silencing that causes *ptsG* gene repression. Further, the coding region of the *ptsG* gene also had 2 SNPs – G13G and A7S. G13G is a synonymous mutation that did not alter the protein sequence whereas A7S is a non-synonymous mutation that altered the protein sequence by changing an alanine base to serine. Further studies need to be done to address this mutation on *ptsG* gene expression.

Apart from these mutations in the promoter and coding region of *ptsG* gene, one of the EvoCrp showed a mutation in the coding region of *avtA* gene that codes for valine-pyruvate aminotransferase. Additionally, 4 of the 5 EvoCrp strains had a mutation in the promoter sequence of *glpF*, a glycerol transporter that carries out facilitated diffusion of glycerol across the inner cell membrane of *E. coli*. Also, all the evolved strains showed a mutation in the intergenic region between the genes *gltP* and *yjcO* which are glutamate/aspartate symporter and putative ABC transporter respectively. The repeated occurrence of mutations in the exact same region of the genome across independently evolved populations reflected the highly convergent nature of adaptive evolution.

To analyze the contribution of the *ptsG* promoter site mutations towards growth recovery, validation of mutations was carried out by site-directed mutagenesis. However, to limit the scale of validation experiment, only one of the mutations in promoter region of *ptsG* gene seen in the strain EvoCrp1, i.e., the mutation IG116 in *ptsG* promoter was validated. This mutation was introduced into a Δ*crp* strain. Growth rate and glucose uptake rate analysis of this strain IG116 Δ*crp*, WT and Δ*crp* were done in shake flask with M9 + 40 mM MOPS + 2 g/L glucose. Growth rate showed a ~85% recovery compared to that of the WT strain (supplementary fig. S5A and S5B). Also, the glucose uptake rate was found to be 8.1±0.1 mM/gDCW/h in IG116 Δ*crp*, 9.2±0.13 mM/gDCW/h in WT and 4.25±0.25 mM/gDCW/h Δ*crp*. It thereby indicated restoration of glucose uptake rate in IG116 Δ*crp* by ~88% to the WT glucose uptake rate.

### Supplementary figures

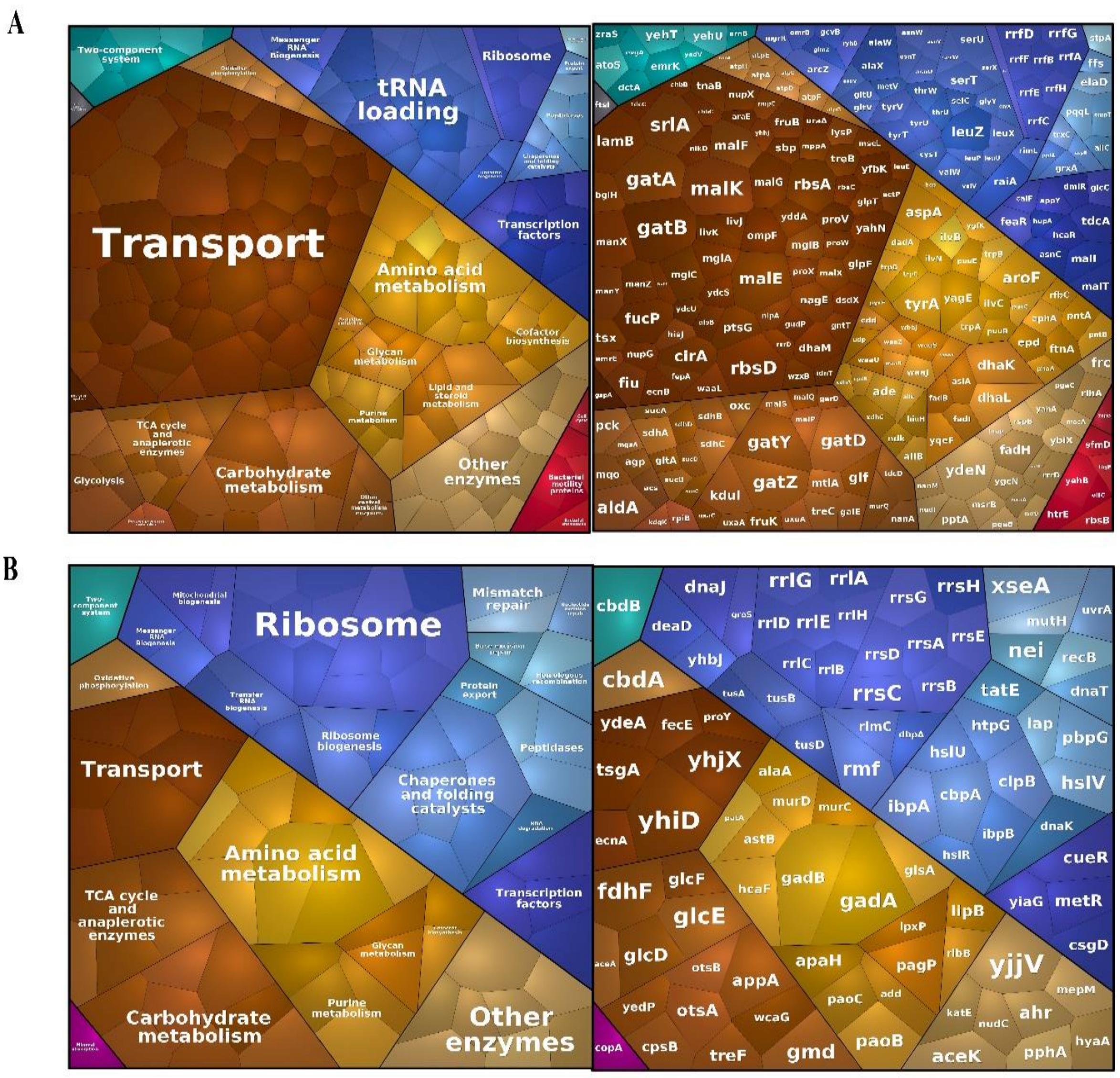


**Supplementary Figure S1.** (A) The Voronoi maps showing the downregulated metabolic pathways and the genes within each pathway, enriched by KEGG classification in Δ*crp*. Transport (adj-P < 0.05), tRNA loading (adj-P < $10^{-4}$) and TCA cycle (adj-P < 0.05) were found to be significantly downregulated. (B) The Voronoi maps showing the upregulated metabolic pathways and the genes within each pathway, enriched by KEGG classification. Ribosome (adj-P < $10^{-5}$), Chaperone and folding catalysts (adj-P < $10^{-2}$) and TCA cycle (adj-P < 0.05) were found to be significantly upregulated. Note: The size of the hexagon within each pathway is directly proportional to the absolute fold change observed for the genes. The color of the hexagon denotes the specific pathways classified by KEGG.

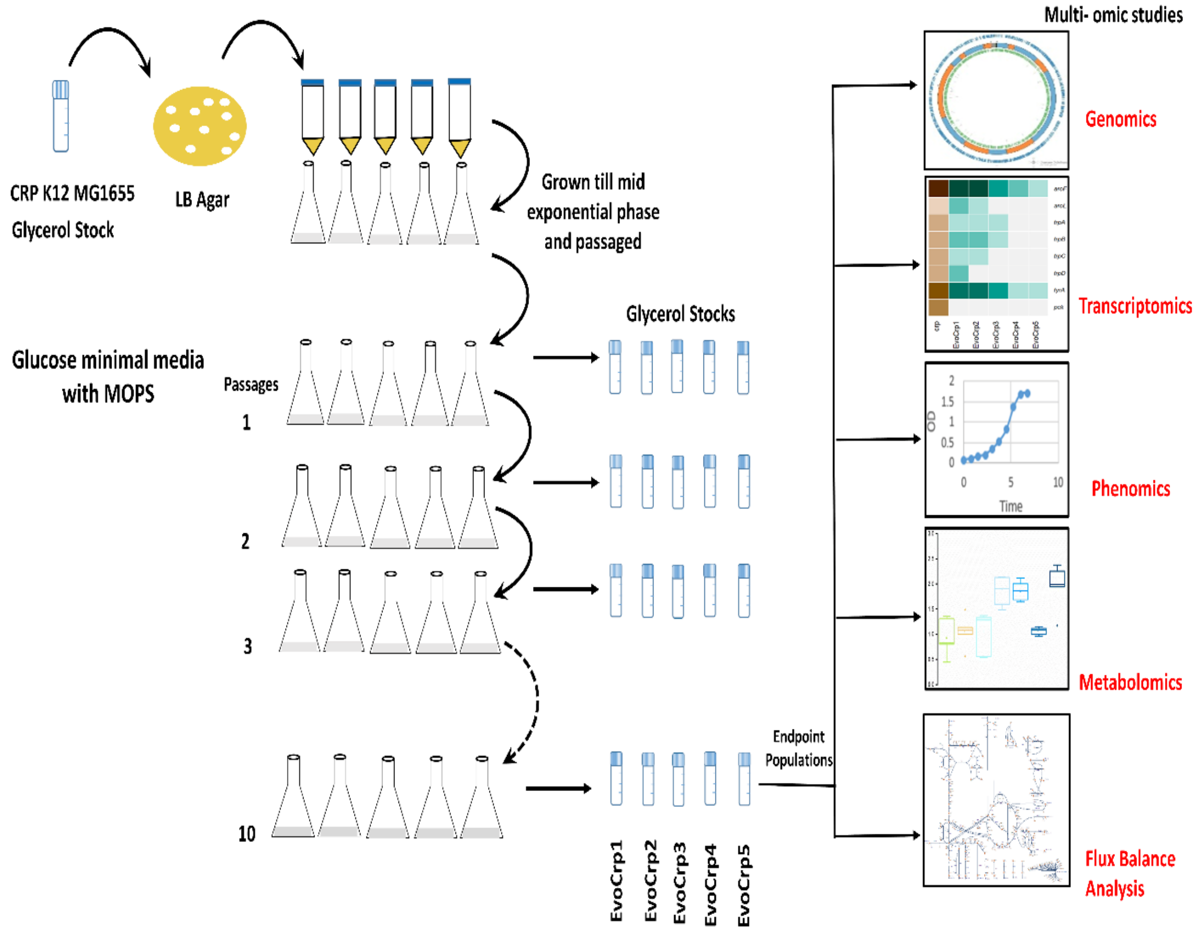


**Supplementary Figure S2.** Overview of the Adaptive Laboratory Experiment (ALE). The figure shows the protocol followed for ALE experiment. Evolution was carried out for 5 replicates of Δ*crp* by passages in M9 + MOPS media with glucose during the mid-exponential phase (0.6-0.7 OD) until a stable growth rate was reached. Glycerol stocks were stored during each passage and PCR checked at each stage to circumvent any contaminations. The end-point population samples were used for further genotypic and phenotypic characterizations.

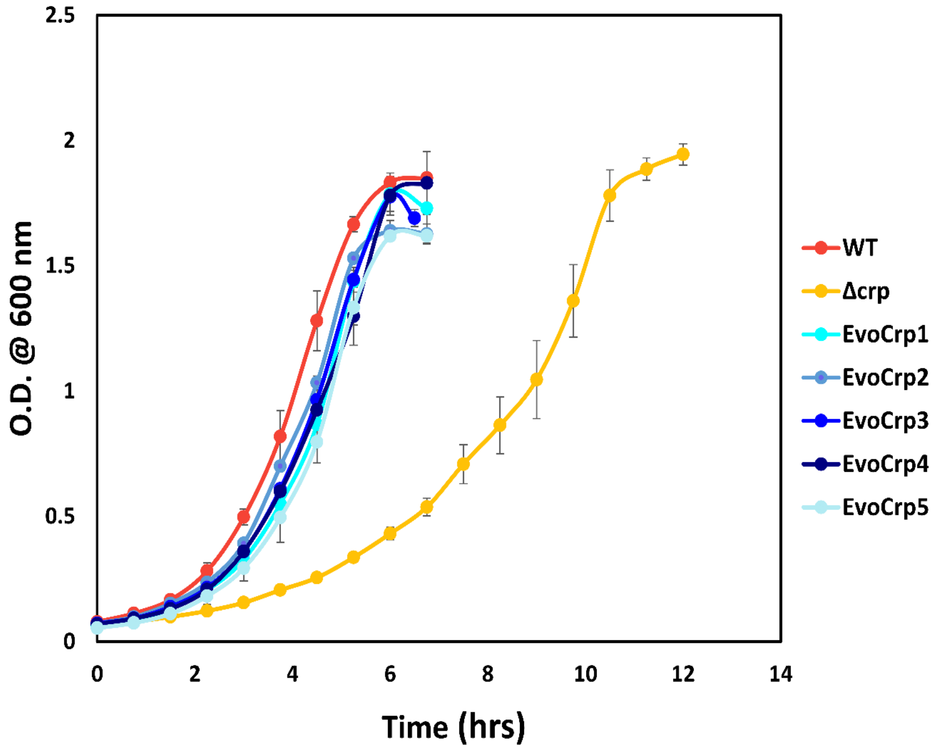


**Supplementary Figure S3.** Growth recovery during evolution of Δ*crp* mutant. Growth curves of the WT, Δ*crp* and endpoint EvoCrp populations- EvoCrp1, EvoCrp2, EvoCrp3, EvoCrp4, and EvoCrp5 which were used for further genotypic and phenotypic characterizations in this study. The curves represent an average of growth curves obtained from 3 biological replicates. The error bars indicate the standard error across the replicates.

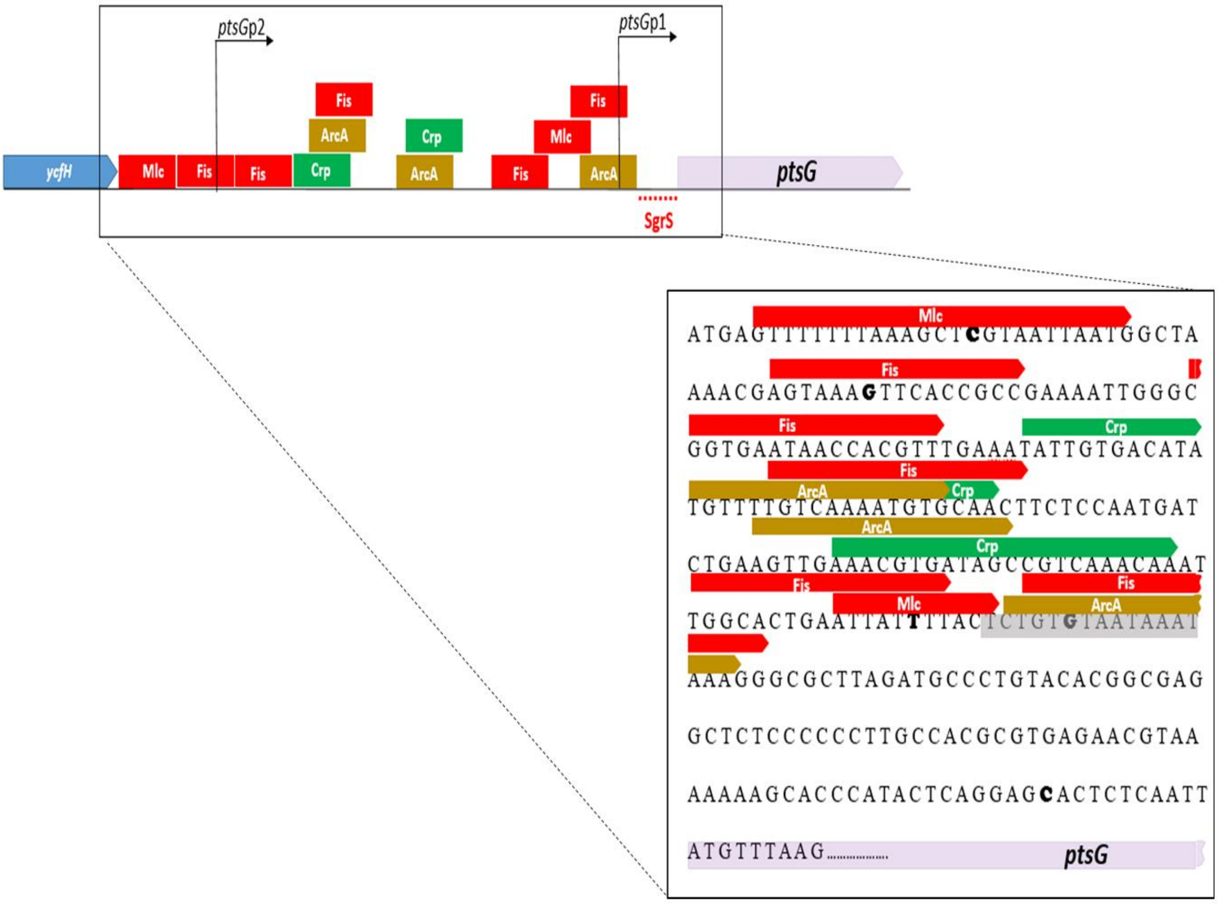


**Supplementary Figure S4.** *ptsG* operon: Depiction on the upstream region of the *ptsG* gene with the positive and negative regulators as reported in EcoCyc. The positive regulators are depicted in green; the negative regulators are depicted in red and dual regulators are depicted in brown. The zoom-in section shows the bases that were mutated in the EvoCrp strains from WGS analysis.

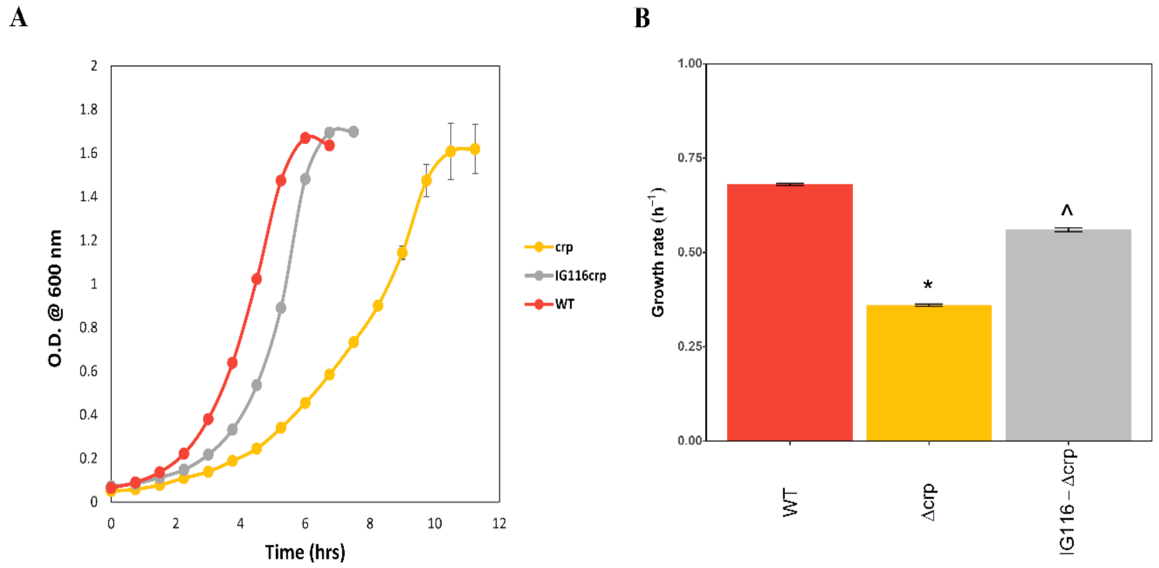


**Supplementary Figure S5.** Growth curves of the WT and the mutants. (A) Growth curves of WT, Δ*crp*, and IG116-Δ*crp* strain generated by *in-vivo* site-directed mutagenesis. The curves represent an average of growth curves obtained from three biological replicates. The error bars indicate the standard error across the replicates. (B) Growth rate recovery caused by IG116-Δ*crp* mutation. The bars represent an average of growth rates obtained from three biological replicates. The error bars indicate the standard error across the replicates. Significance of decrease in Δ*crp* vs WT is shown by asterisk whereas significance of increase in IG116-Δ*crp* vs Δ*crp* is shown by caret.

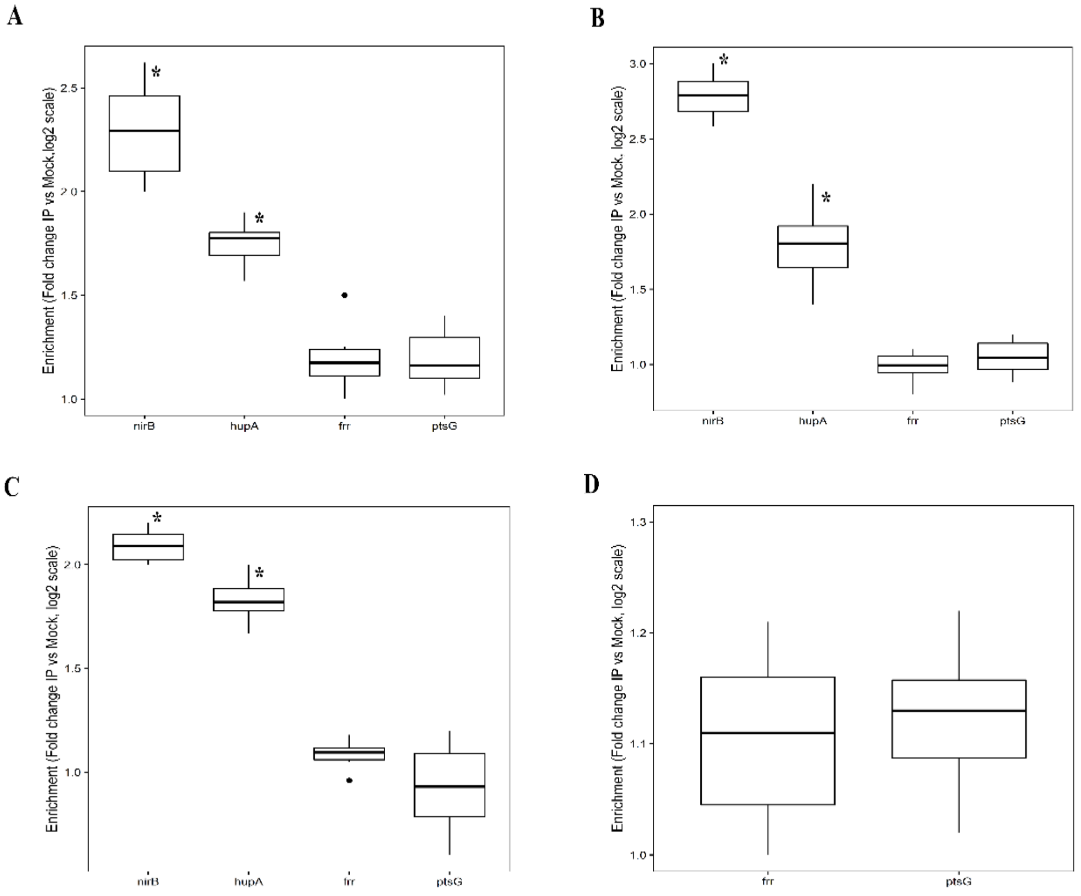


**Supplementary Figure S6.** ChIP qPCR for *ptsG* against immuno-precipitated DNA. (A) Enrichment of *ptsG* IP-DNA vs mock-DNA in WT Fis-FLAG strain with intergenic regions of *nirB* and *hupA* as positive controls and *frr* coding region as the random region. (B) Enrichment of *ptsG* IP-DNA vs mock-DNA in Δ*crp* Fis-FLAG strain with *nirB* and *hupA* as positive controls and *frr* as the random region. (C) Enrichment of *ptsG* IP-DNA vs mock-DNA in IG116-Δ*crp* Fis-FLAG strain with *nirB* and *hupA* as positive controls and *frr* as the random region. (D) Enrichment of *ptsG* IP-DNA vs mock-DNA in IG116-Δ*crp* Mlc-FLAG strain with *frr* as the random region. Fold change was calculated by using $2^{-\Delta\Delta C_t}$ of the immuno-precipitated and mock threshold cycle ($C_t$). Results are shown for the qPCR for two biological and three technical replicates of each sample. Significance was calculated using Wilcoxon rank sum test ($P < 0.01$) compared to the random region (*frr*) and are shown with asterisk.

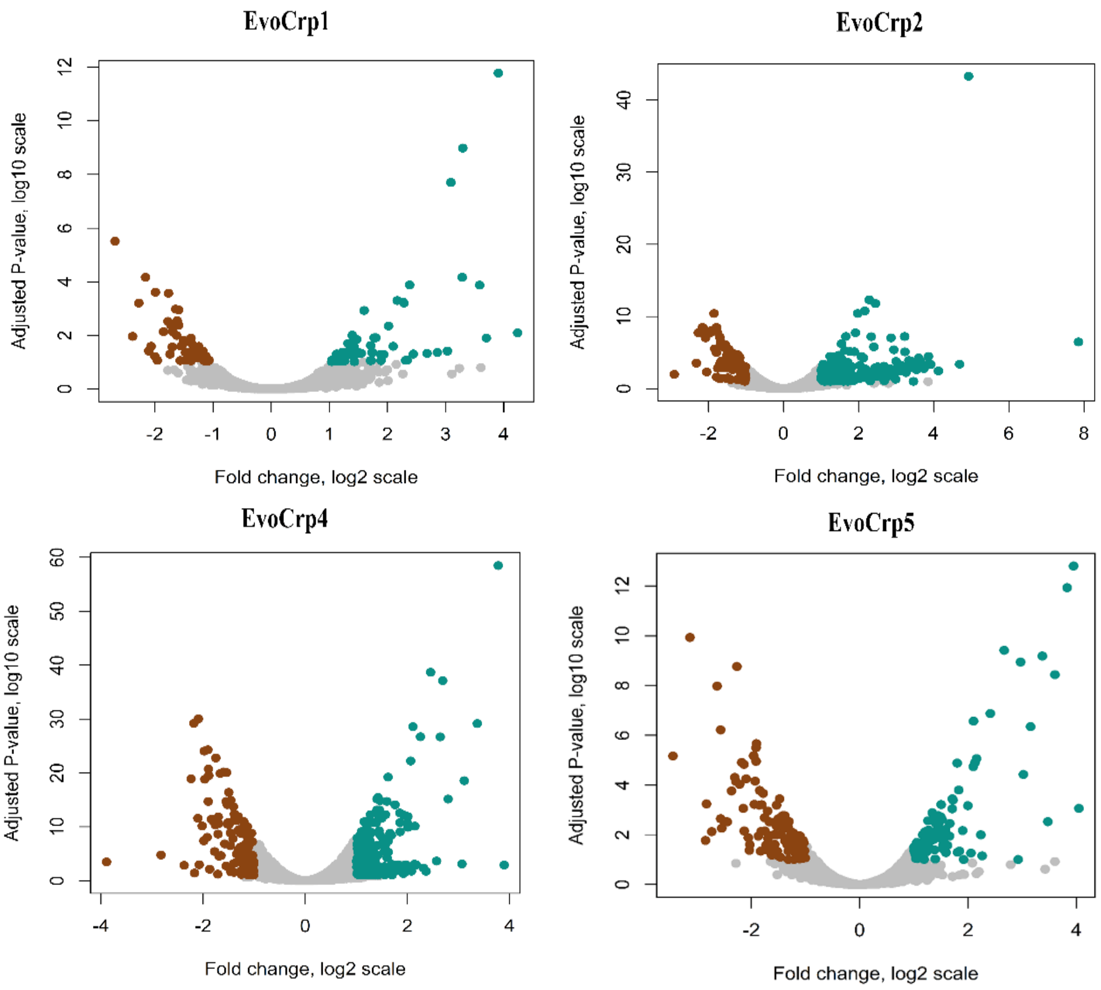


**Supplementary Figure S7.** Volcano plot of the DE genes of EvoCrp1,2,4,5 vs Δ*crp* depicted as adjusted (Adj.) P-value (log10 scale) vs Fold change (log2 scale). The brown dots indicate downregulated genes and the cyan dots indicate upregulated genes.

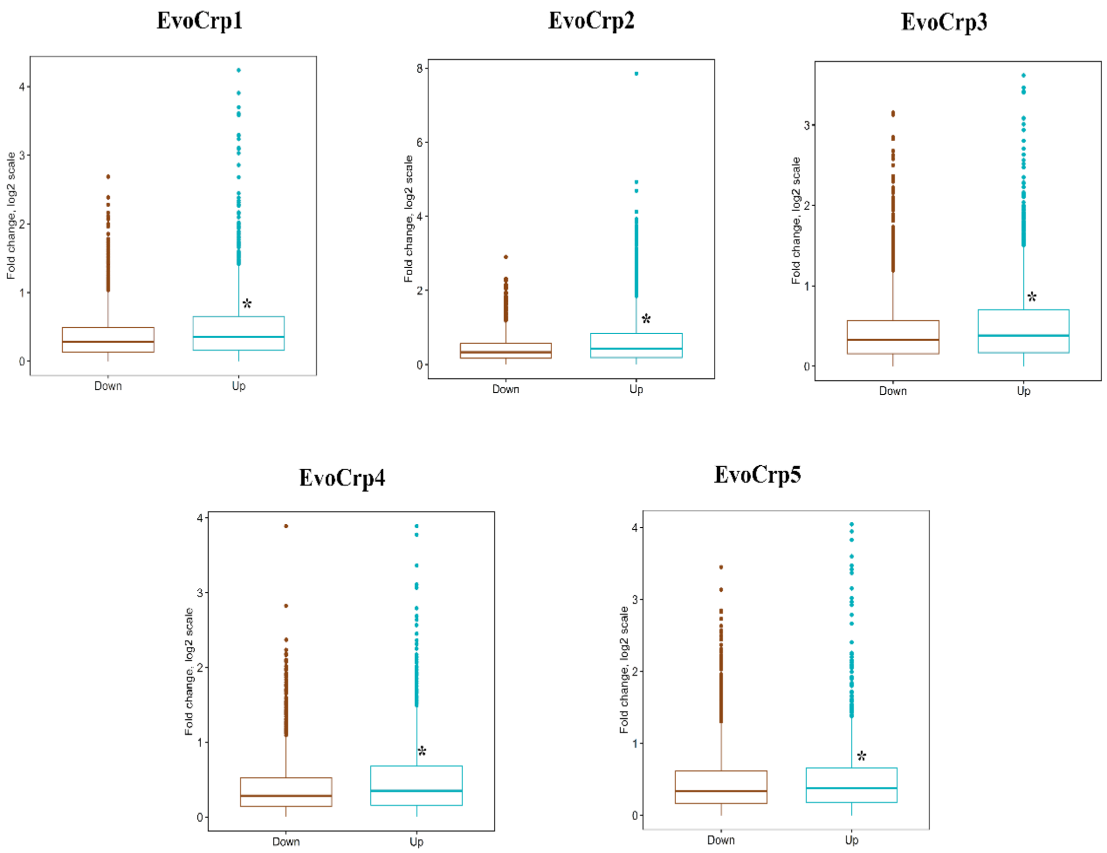


**Supplementary Figure S8.** Box-plots depicting the median fold change (absolute) of all the upregulated genes (cyan) and the downregulated genes (brown) of the EvoCrp compared to Δ*crp* mutant. Significance in median expression levels were calculated using Mann-Whitney test and indicated by asterisk ($P < 10^{-6}$ for all the EvoCrp strains and $P < 0.015$ for EvoCrp5).

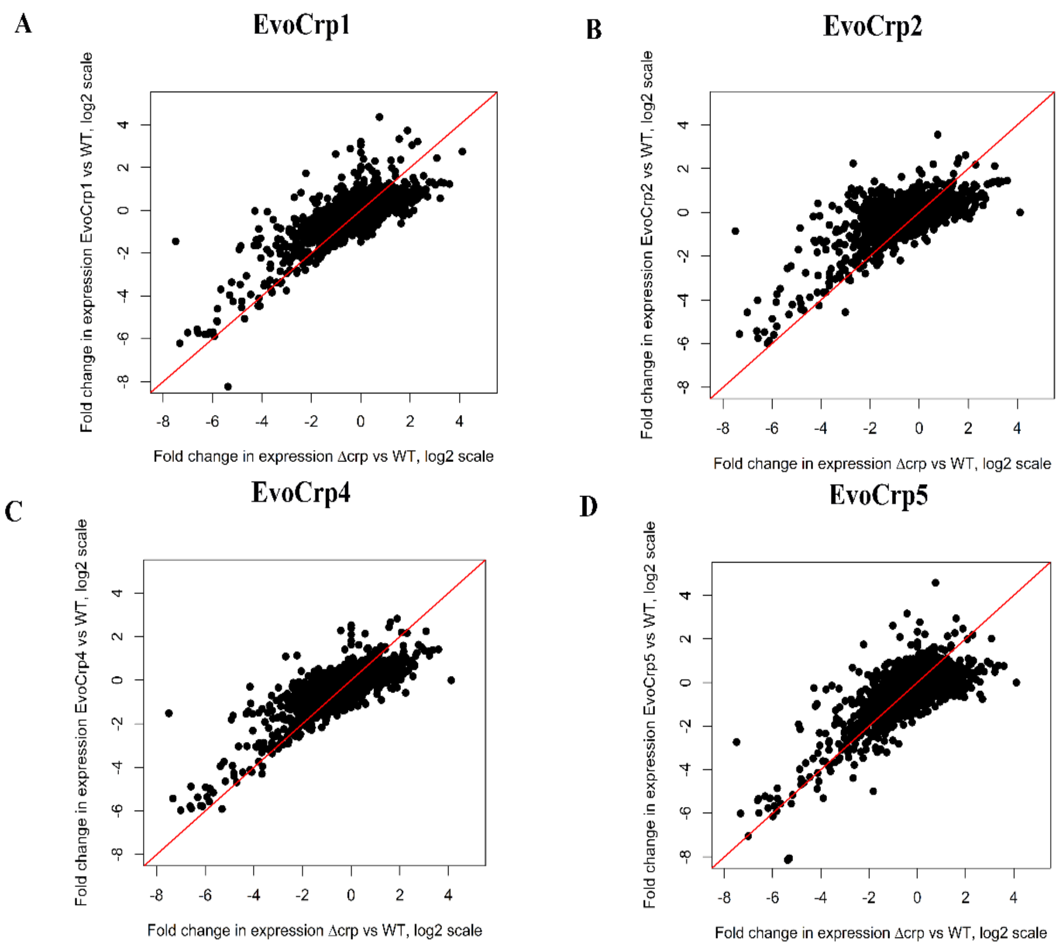


**Supplementary Figure S9.** Correlation plots between EvoCrp vs WT and Δ*crp* vs WT. (A) EvoCrp1 vs WT and Δ*crp* vs WT (Pearson correlation coefficient, r = 0.79, P < $10^{-15}$). (B) EvoCrp2 vs WT and Δ*crp* vs WT. (Pearson correlation coefficient, r = 0.69, P < $10^{-15}$). (C) EvoCrp4 vs WT and Δ*crp* vs WT (Pearson correlation coefficient, r = 0.80, P < $10^{-15}$). (D) EvoCrp5 vs WT and Δ*crp* vs WT (Pearson correlation coefficient, r = 0.76, P < $10^{-15}$).

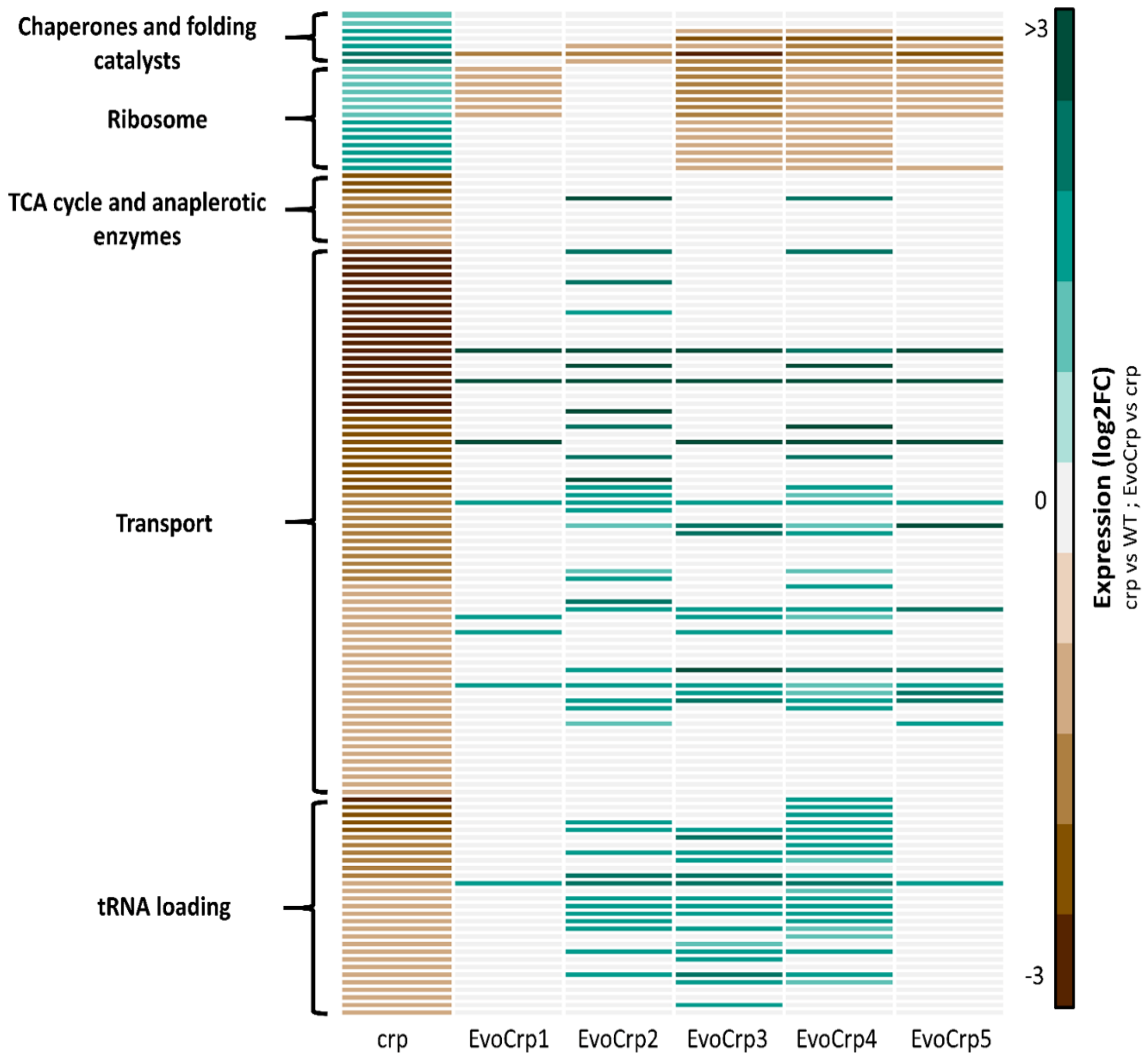


**Supplementary Figure S10.** Heatmap depicting the pathways found to be significantly altered in Δ*crp*. After evolution, these specific pathways were found to be significantly enriched in EvoCrp strains, thereby indicating recovery in the gene expression pattern.

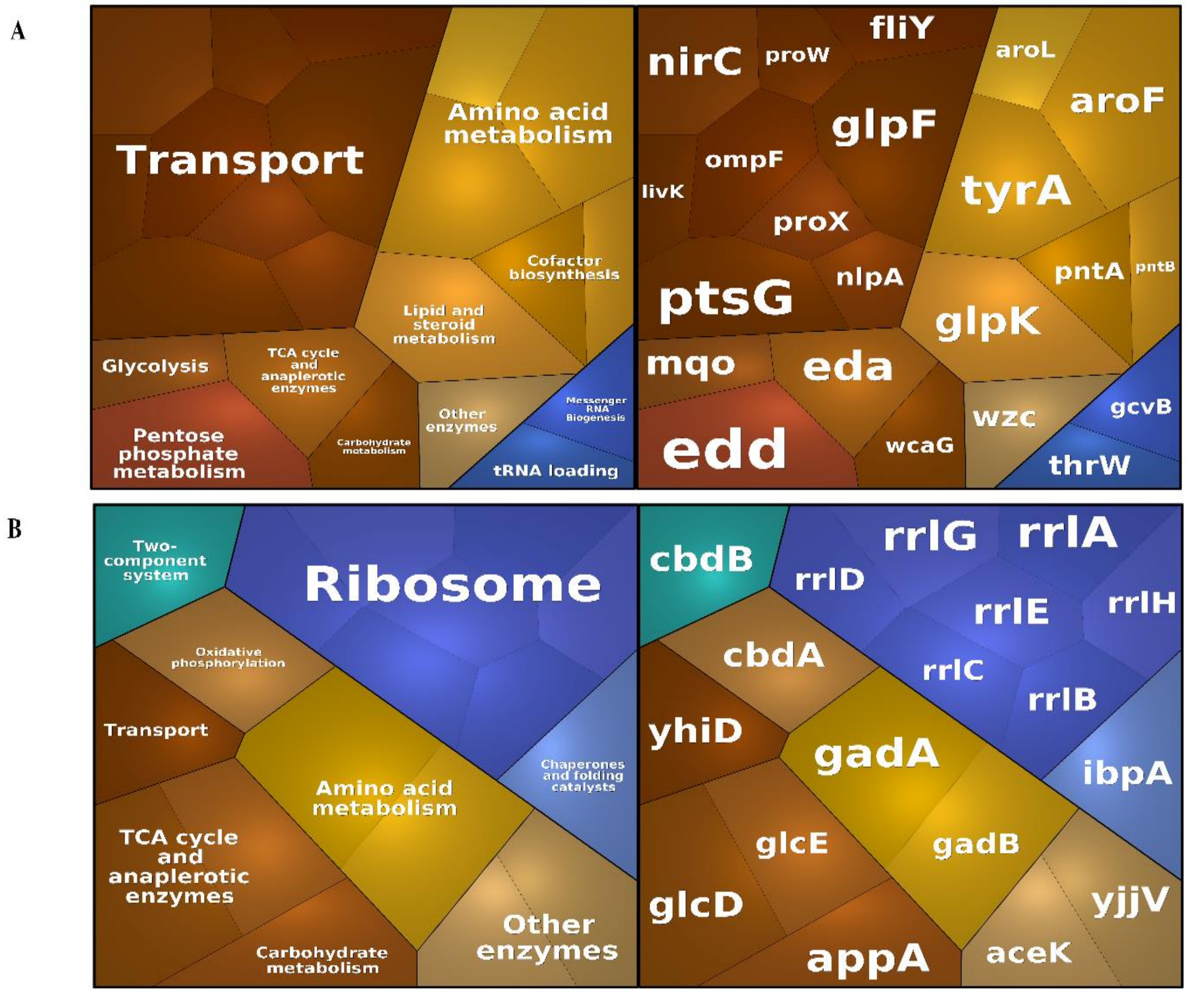


**Supplementary Figure S11.** Enrichment analysis of the DE genes in EvoCrp1 vs Δ*crp* by KEGG classification respectively. (A)The upregulated metabolic pathways and its genes in EvoCrp1. Transport (adj-P < $10^{-2}$) was found to be significantly upregulated. (B) The downregulated metabolic pathways and its genes in EvoCrp1. Ribosome (adj-P < $10^{-5}$) was found to be significantly downregulated.

A

B

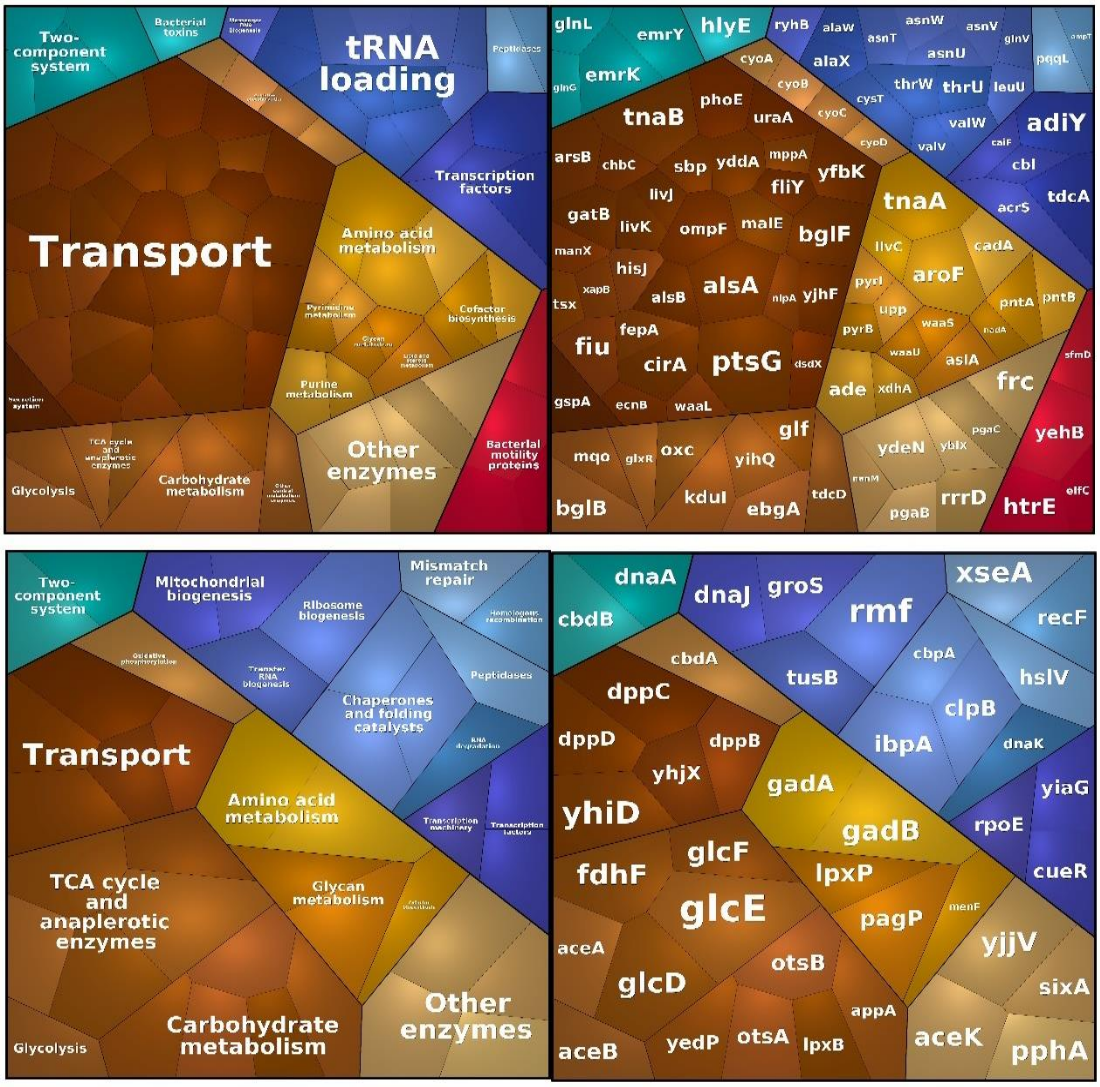


**Supplementary Figure S12.** Enrichment analysis of the DE genes in EvoCrp2 vs Δ*crp* by KEGG classification respectively. (A)The upregulated metabolic pathways and its genes in EvoCrp2. tRNA loading (adj-P < $10^{-2}$) was found to be significantly upregulated. (B) The downregulated metabolic pathways and its genes in EvoCrp2. No pathways were found to be significantly downregulated.

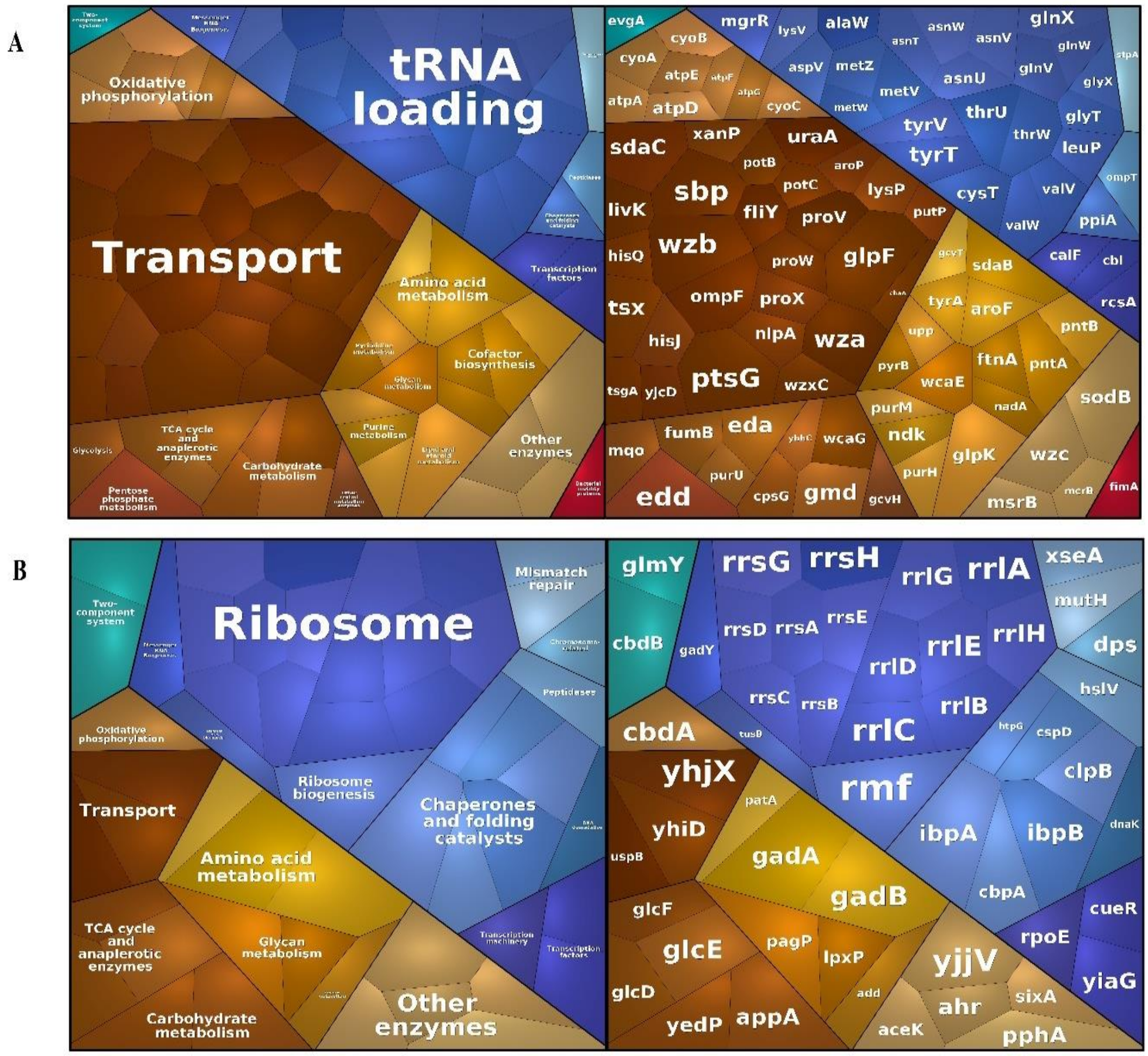


**Supplementary Figure S13.** Enrichment analysis of the DE genes in EvoCrp3 vs Δ*crp* by KEGG classification respectively. (A)The upregulated metabolic pathways and its genes in EvoCrp3. Transport (adj-P < $10^{-3}$), tRNA loading (adj-P < $10^{-13}$) and Oxidative phosphorylation (adj-P < $10^{-5}$) were found to be significantly upregulated. (B) The downregulated metabolic pathways and its genes in EvoCrp3. Ribosome (adj-P < $10^{-8}$), and Chaperone and folding catalysts (adj-P < $10^{-2}$) were found to be significantly downregulated.

A

B

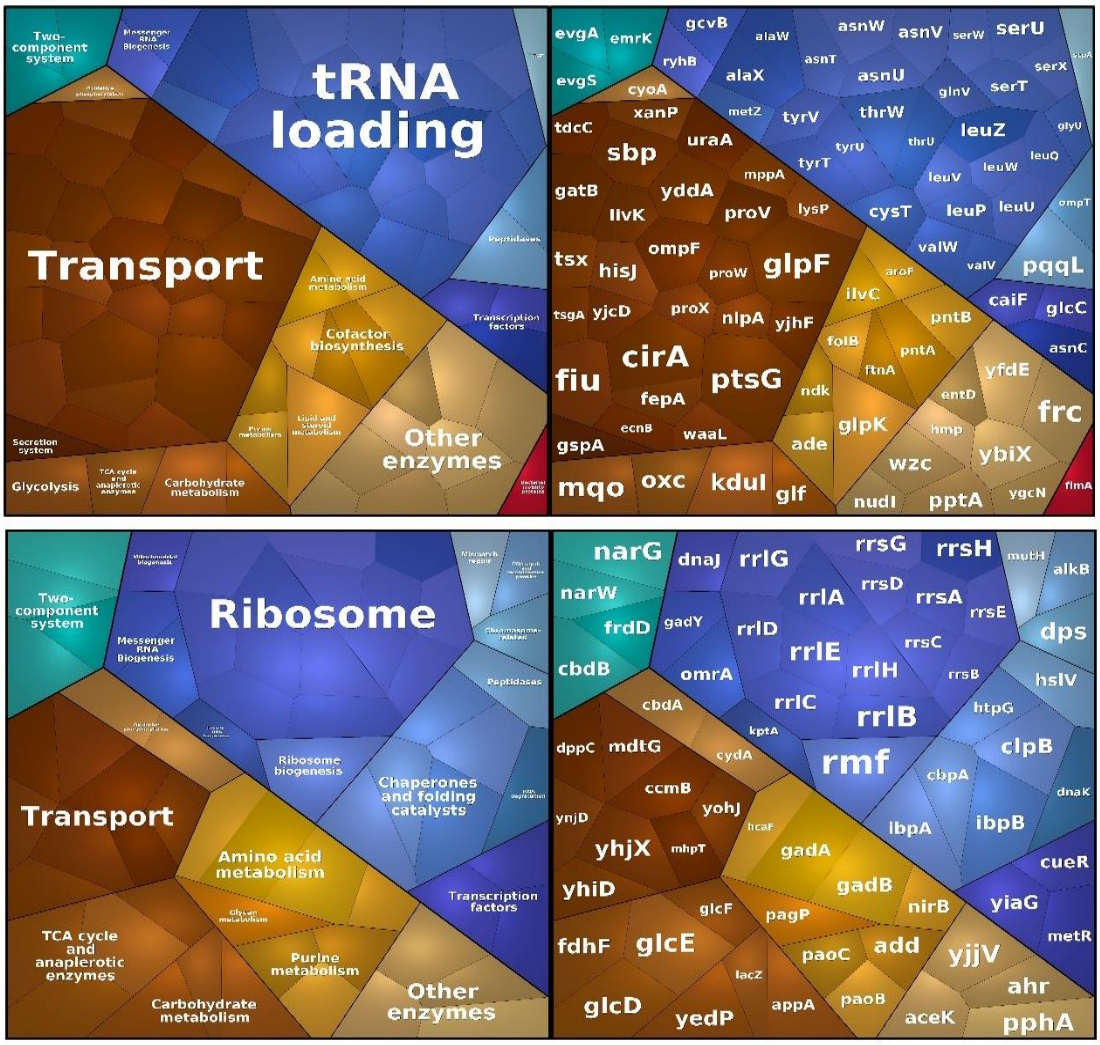


**Supplementary Figure S14.** Enrichment analysis of the DE genes in EvoCrp4 vs Δ*crp* by KEGG classification respectively. (A)The upregulated metabolic pathways and its genes in EvoCrp4. Transport (adj-P < 0.05), and tRNA loading (adj-P < $10^{-15}$) were found to be significantly upregulated. (B) The downregulated metabolic pathways and its genes in EvoCrp4. Ribosomes (adj-P < $10^{-17}$), and Chaperones and folding catalysts (adj-P < 0.05) were found to be significantly downregulated.

A

B

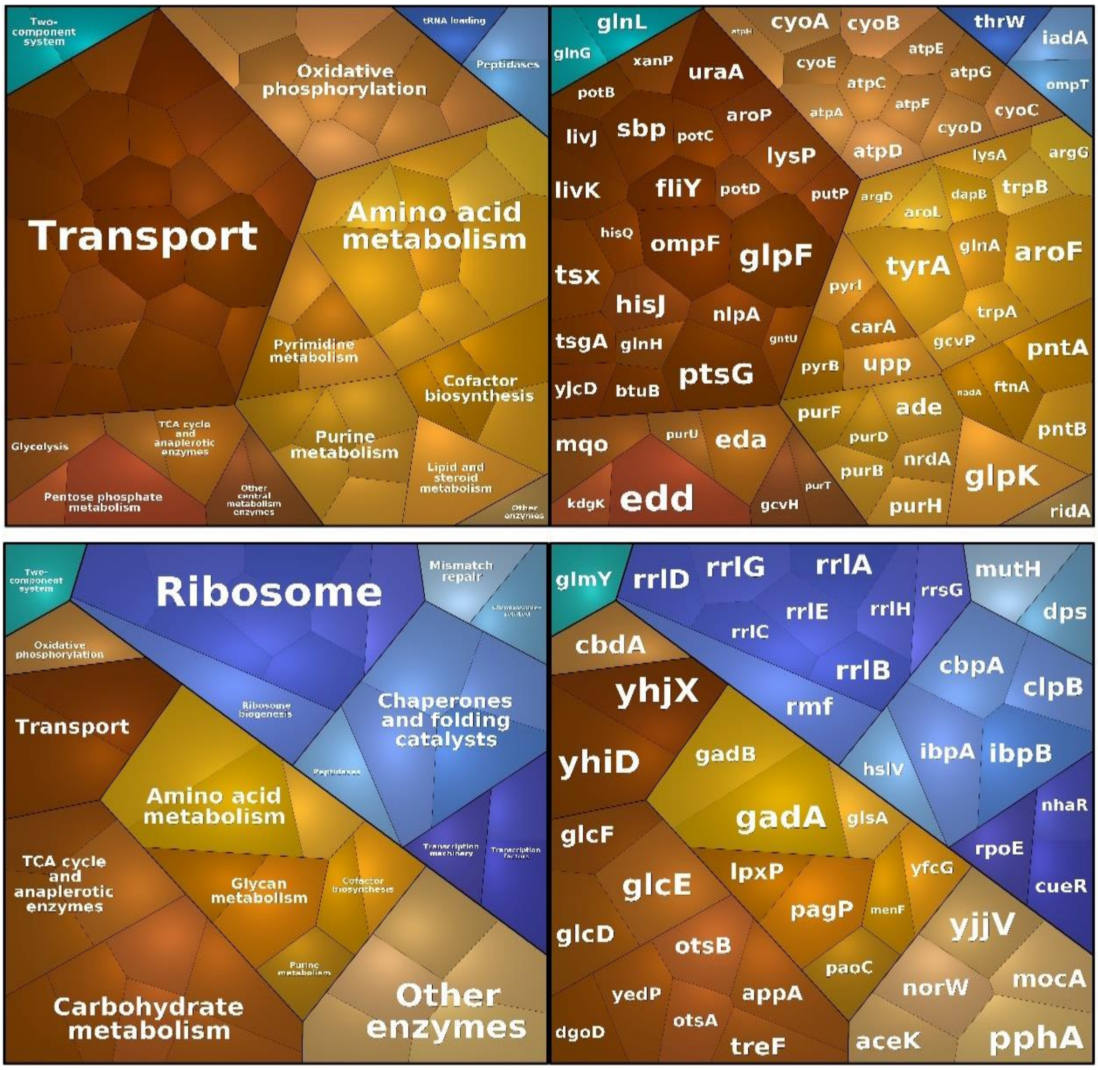


**Supplementary Figure S15.** Enrichment analysis of the DE genes in EvoCrp5 vs Δ*crp* by KEGG classification respectively. (A)The upregulated metabolic pathways and its genes in EvoCrp5. Transport (adj-P < $10^{-4}$), Amino acid metabolism (adj-P < $10^{-3}$), Oxidative phosphorylation (adj-P < $10^{-12}$) and Purine metabolism (adj-P < $10^{-3}$) were found to be significantly upregulated. (B) The downregulated metabolic pathways and its genes in EvoCrp5. Carbohydrate metabolism (adj-P <0.05), and Ribosomes (adj-P < $10^{-2}$) were found to be significantly downregulated.

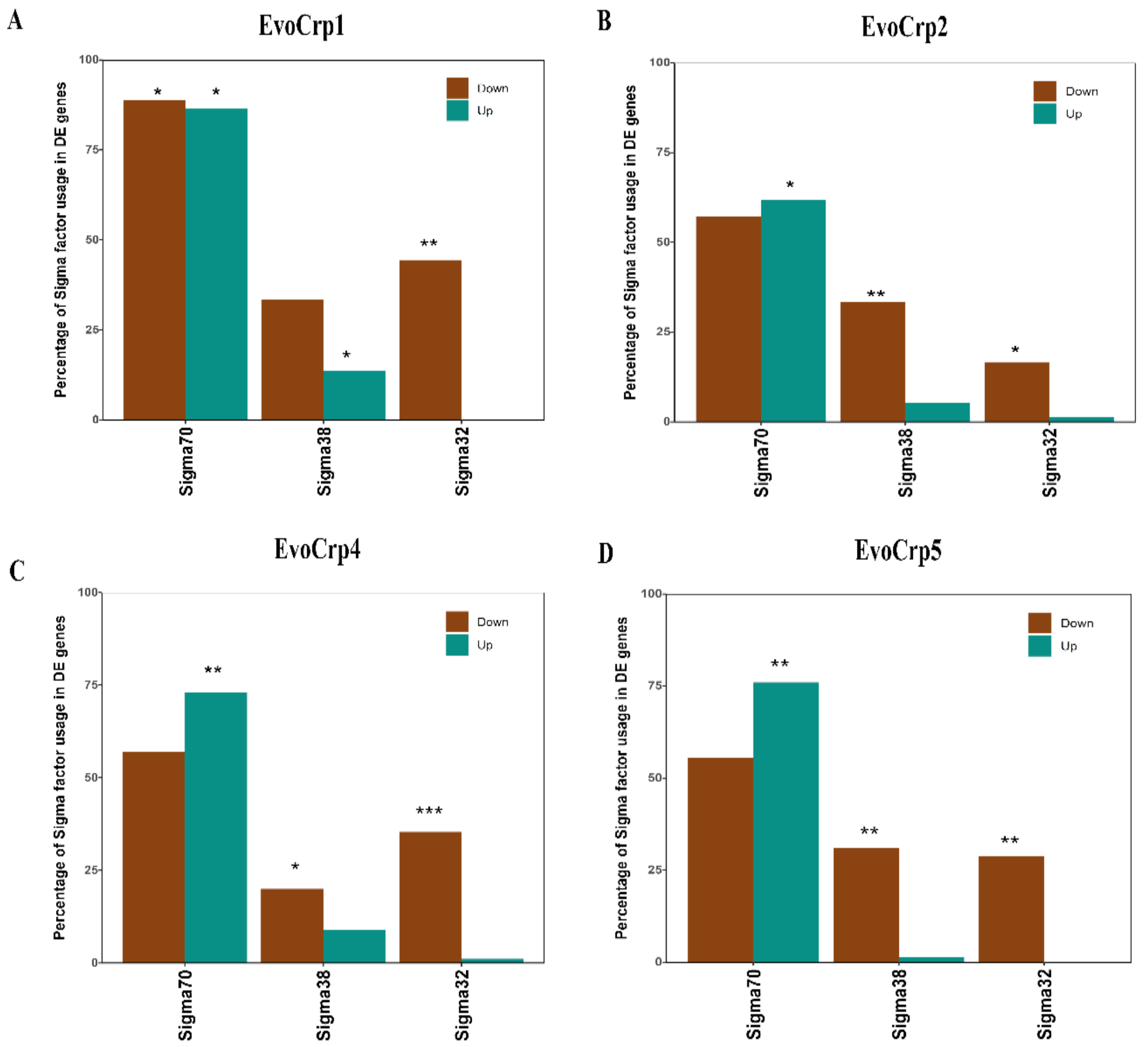


**Supplementary Figure S16.** Enrichment of sigma factors on metabolic pathway related genes in evolved populations. The brown bars and the cyan bars indicate the percentage of downregulated and upregulated genes in EvoCrp strains vs Δ*crp* respectively. Significant increase or decrease is denoted by asterisks: one asterisk indicates $P < 0.05$, two asterisks indicate $P < 10^{-4}$ and three asterisks indicate $P < 10^{-10}$.

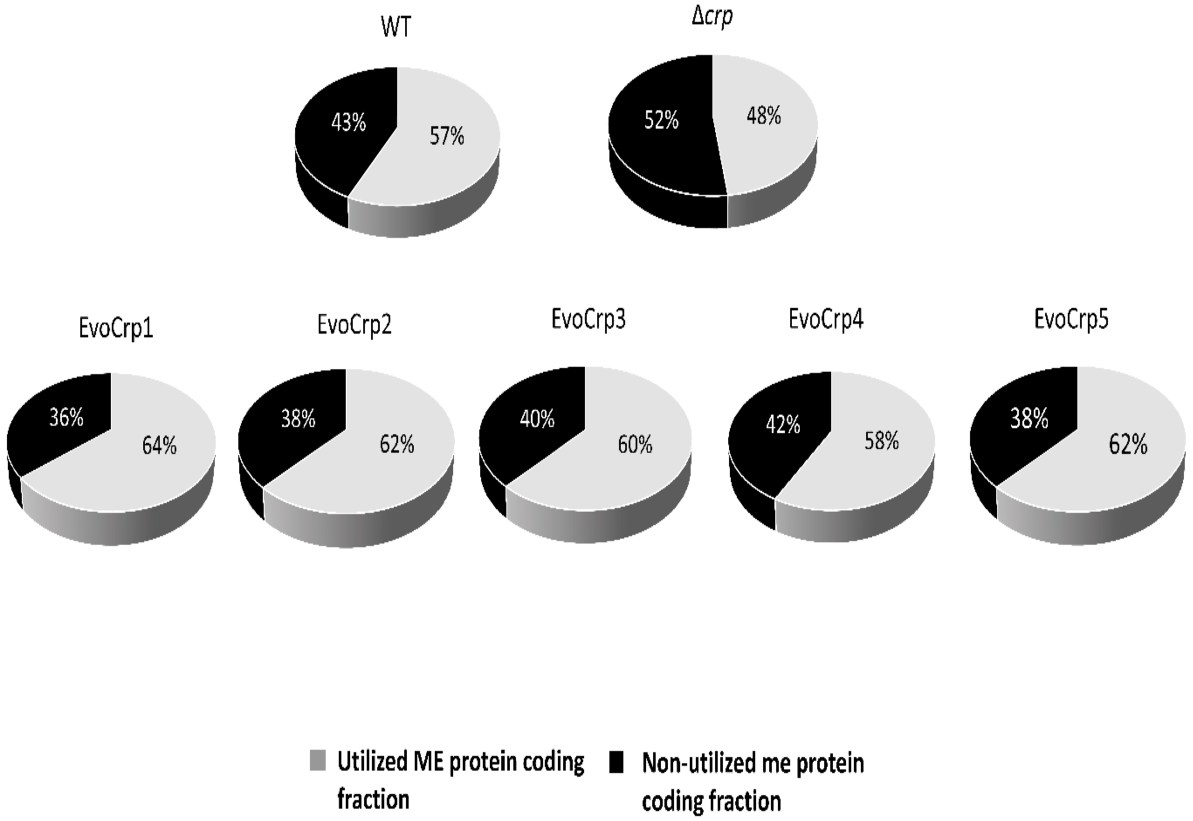


**Supplementary Figure S17.** Changes observed in the percentage of proteome allocation towards uME (grey) and nonME (black) in WT, Δ*crp* and EvoCrp strains depicted as pie-chart.

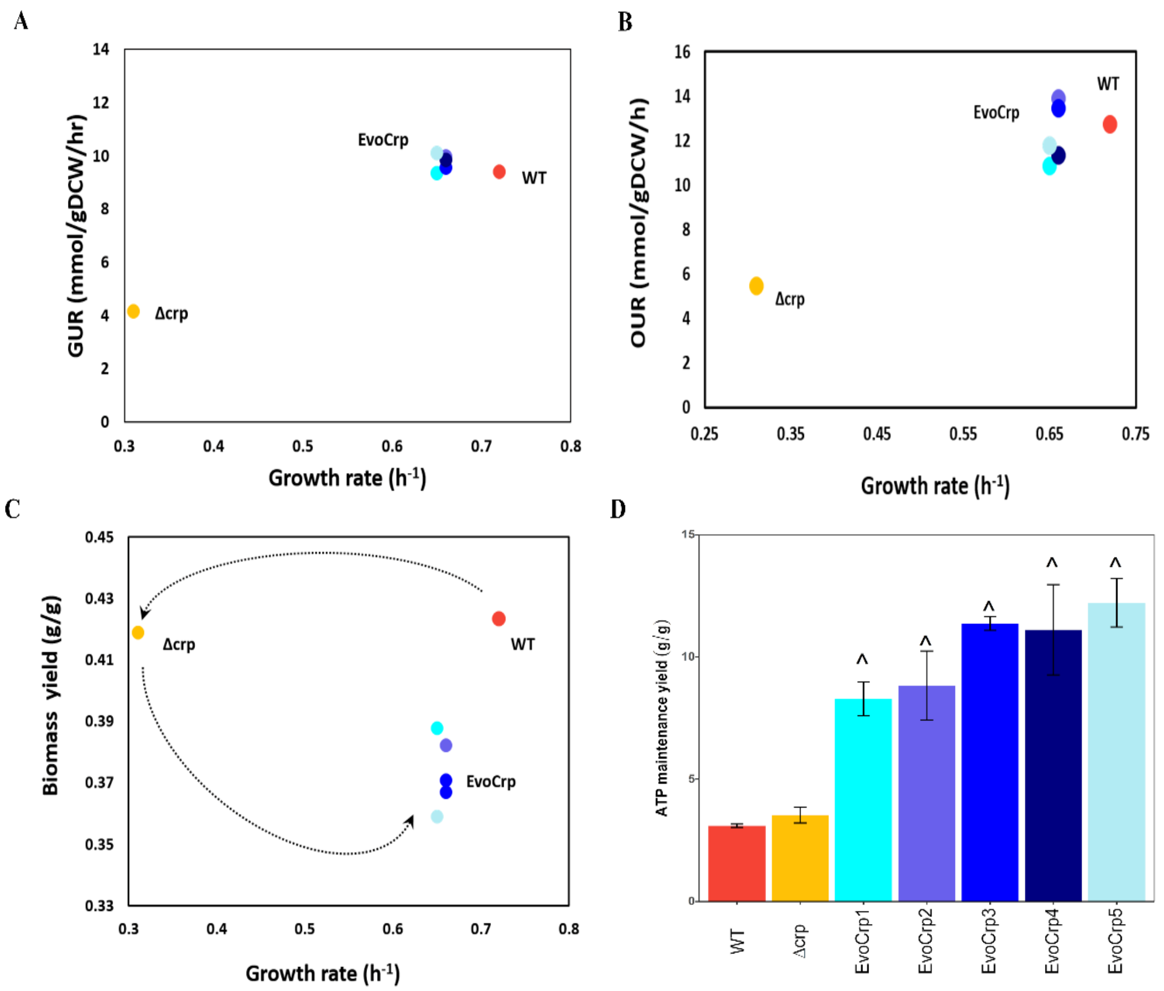


**Supplementary Figure S18.** (A) Pearson pairwise correlation between growth rates and glucose uptake rates (GUR) of all the strains (Pearson correlation coefficient, r = 0.96, P < $10^{-3}$). (B) Pearson pairwise correlation between growth rates and oxygen uptake rates (OUR) of all the strains (Pearson correlation coefficient, r = 0.92, P < $10^{-2}$). (C) Pearson pairwise correlation between growth rates and biomass yields (Pearson correlation coefficient, r = -0.87, P < 0.05, for EvoCrp and *Δcrp*). The dotted curves show the separation of the Δ*crp* and WT and migration of EvoCrp back towards WT as a result of evolution. (D) Bar-plot depicting ATP maintenance (ATPM) yields expressed as (g/g) was predicted from flux balance analysis for WT, Δ*crp* and the EvoCrp strains using ATPM maximization as the objective function. The bars represent an average of rate and yields obtained from three biological replicates. The error bars indicate the standard error across the replicates. Significance of decrease in EvoCrp vs Δ*crp* is shown by caret.

| **Strain Name** | **Description** | **Source** |
|---|---|---|
| WT | *E. coli* K12 MG1655 | Keio collection (CGSC #6300) |
| Δ*crp* | *E. coli* K12 MG1655 with *crp* gene deletion | This work |
| EvoCrp1 | Population 1 of Δ*crp* evolved for ~100 generation | This work |
| EvoCrp2 | Population 2 of Δ*crp* evolved for ~100 generation | This work |
| EvoCrp3 | Population 3 of Δ*crp* evolved for ~100 generation | This work |
| EvoCrp4 | Population 4 of Δ*crp* evolved for ~100 generation | This work |
| EvoCrp5 | Population 5 of Δ*crp* evolved for ~100 generation | This work |
| IG116-Δ*crp* | Δ*crp* strain with an intergenic mutation 116 bp upstream of translation start site | This work |
| WT-Fis- FLAG | WT with Fis-FLAG tag | This work |
| Δ*crp*-Fis-FLAG | Δ*crp* with Fis-FLAG tag | This work |
| IG116-Δ*crp*-Fis-FLAG | IG116-Δ*crp* with Fis-FLAG tag | This work |
| IG116-Δ*crp*-Mlc-FLAG | IG116-Δ*crp* with Mlc-FLAG tag | This work |

**Supplementary Table S1.** Strains used in this study

| Gene | Protein change | Mutation | Genome Position | Annotation (this study) | EvoCrp1 | EvoCrp2 | EvoCrp3 | EvoCrp4 | EvoCrp5 |
|---|---|---|---|---|---|---|---|---|---|
| *ptsG* promoter p2 | Intergenic | C → T | 1157593 | IG276 | 26.9% | | 13.9% | 11% | 15.9% |
| *ptsG* promoter p1 | Intergenic | T → G | 1157753 | IG116 | 72.6% | | | | |
| *ptsG* promoter p1 | Intergenic | (TCTGTGTAATAAAT)$_{1\rightarrow 2}$ | 1157771 | IG98 | | | | | 69.4% |
| *ptsG* gene | Coding | T → A | 1157907 | G13G | | | | | 5.5% |
| *ptsG* promoter p1 | Intergenic | G → A | 1157763 | IG106 | | 60.7% | | 9.3% | |
| Upstream of *ptsG* gene | Intergenic | C → A | 1157858 | IG11 | | 32.3% | | | |
| *ptsG* gene | Coding | G → T | 1157887 | A7S | | | 78.5% | | |
| *ptsG* promoter p2 | Intergenic | G → A | 1157619 | IG250 | | | | 80.5% | |
| *glpF* promoter | Intergenic | +4 bp | 4118284 | IG194 | 71.2% | 68.2% | | 67.3% | 67.5% |
| *gltP* /yjcO | Intergenic | C → T | 4296060 | | 27.6% | 27.3% | 30.3% | 29.5% | 28.6% |
| avtA *gene* | Coding | C → T | 3740019 | G105G | | | 11% | | |

**Supplementary Table S2:** List of mutations identified across the different EvoCrp strains. Lane 1: Gene description, Lane 2: The intragenic or intergenic nature of mutation, Lane 3: The type of mutations identified (mostly SNPs with 1 duplication and 1 insertion), Lane 4: Genomic coordinates of the identified mutation, Lane 5: Annotation for the mutations (Note: In case of intergenic mutations, IG denotes intergenic and the numbers following IG denotes the bases upstream of the translation start site where the mutation was detected. In cases of coding region mutation, the annotation denotes the amino acid change in the protein sequence), Lane 6- 10: The mutation frequency (in percentage) of the mutation within the replicate populations.

| **Strains** | **Growth rate ($h^{-1}$)** | **Biomass yield (g/g)** | **Specific rates(mM/gDCW/h)** | | |
|---|---|---|---|---|---|
| | | | **Glucose** | **Acetate** | **Oxygen** |
| **WT** | 0.72 ± 0.01 | 0.43 ± 0 | 9.4 ± 0.18 | 6.62 ± 0.38 | 12.75 ± 0.18 |
| **Δ*crp*** | 0.31 ± 0.01 | 0.42 ± 0.01 | 4.15 ± 0.09 | 2.91 ± 0.05 | 5.45 ± 0.14 |
| **EvoCrp1** | 0.65 ± 0.01 | 0.39 ± 0.01 | 9.34 ± 0.19 | 6.16 ± 0.14 | 10.877 ± 0.57 |
| **EvoCrp2** | 0.66 ± 0.01 | 0.37 ± 0.01 | 9.55 ± 0.23 | 6.46 ± 0.23 | 13.91 ± 0.54 |
| **EvoCrp3** | 0.66 ± 0.01 | 0.37 ± 0.02 | 9.99 ± 0.2 | 6.29 ± 0.1 | 13.44 ± 0.07 |
| **EvoCrp4** | 0.66 ± 0.007 | 0.38 ± 0 | 9.86 ± 0.39 | 6.09 ± 0.26 | 11.33 ± 0.19 |
| **EvoCrp5** | 0.65 ± 0.01 | 0.37 ± 0 | 10.11 ± 0.28 | 6.41 ± 0.02 | 11.78 ± 0.52 |

**Supplementary Table S3.** Physiological characterization of WT, Δ*crp* and EvoCrp strains. The measurements indicate the average of three biological replicates. The error bars indicate the standard error across the replicates.
